# Peripheral NOD-like receptor deficient inflammatory macrophages trigger neutrophil infiltration disrupting daytime locomotion

**DOI:** 10.1101/2021.10.27.466033

**Authors:** Victoria Kwon, Peiwen Cai, Cameron T. Dixon, Victoria Hamlin, Caroline G. Spencer, Alison M. Rojas, Celia E. Shiau

## Abstract

Inflammation is known to disrupt normal behavior, yet the underlying neuroimmune interactions remain elusive. Here, we investigated whether inappropriate macrophage-evoked inflammation alters CNS control of daily-life animal locomotion using a set of zebrafish mutants selected for specific macrophage dysfunction and microglia deficiency. Large-scale genetic and computational analyses revealed that NOD-like receptor *nlrc3l* mutants are capable of normal motility and visuomotor response, but preferentially swim less in the daytime, suggesting low motivation rather than physical impairment. Examining their brain activities and structures implicate impaired dopaminergic descending circuits, where neutrophils abnormally infiltrate. Furthermore, neutrophil depletion recovered daytime locomotion. Restoring wild-type macrophages reversed behavioral and neutrophil aberrations, while three other microglia-lacking mutants failed to phenocopy *nlrc3l* mutants. Overall, we reveal how peripheral inflammatory macrophages with elevated pro-inflammatory cues (including *il1b*, *tnfa*, *cxcl8a*) in the absence of microglia co-opt neutrophils to infiltrate the brain, thereby enabling local modulation of neural circuits affecting spontaneous locomotion.

## Introduction

The impact of the immune system on brain function is seemingly self-evident when recognizing that behavioral changes coined as “sickness behaviors” are triggered by the body’s response to infection. These are a collection of behaviors evolutionarily conserved from zebrafish to humans, which include fatigue, social withdrawal, and reduced physical activity intended to conserve energy to meet the demands of fighting off pathogens for survival^1–5^. In response to an infection or injury, immune cells collectively instigate inflammation that serves two main purposes: to eliminate pathogens or harmful debris and restore homeostasis^6, 7^. To this end, innate immune cells are thought to release pro-inflammatory cytokines predominantly IL-1β and TNF-alpha in the periphery that subsequently can reach the brain parenchyma via passage through the extracellular fluids or through the peripheral nerves^3, 8–11^. The brain is thought to be able to recognize these pro-inflammatory cytokines as cues to implement the sickness behavior circuit^2, 9^. Besides infection, immune dysregulation can also lead to unwarranted inflammation causing high elevation of these cytokines systemically^6^, but whether this can cause sickness behavior remains unclear. Irrespective of the infection context, the underlying mechanisms by which inflammation triggers behavioral changes remain poorly understood, including the dynamic neuroimmune interactions and brain targets involved.

Macrophages are key players that orchestrate inflammation to promote tissue resolution after an infection, injury or disease ^12–15^, but how their dysregulation in the periphery may alter brain function and its behavioral outcome remains underexplored. A recent study indicated that peripheral macrophages along the peripheral nerve were responsive to spinal cord neurodegeneration and able to repress proinflammatory microglial activation^16^, implicating more intimate communication between peripheral macrophages and central nervous system (CNS) cells than previously thought. In terms of routes through which peripheral immune signals may be interpreted by the brain, microglia, brain-resident macrophages, may act as major CNS facilitators for such communications as they can respond rapidly to environmental changes^17^. Furthermore, intimate ties between peripheral adaptive immunity and CNS functions have been well demonstrated through active engagement of T-cells in the brain meninges at homeostasis and in disease states capable of disrupting CNS circuits that affect animal behaviors^18–22^. Therefore, the interplay of the peripheral immune system, both the innate and adaptive arms, and the CNS is emerging as an important factor shaping brain function and homeostasis.

Genetic basis for immune crosstalk on neural circuits and behavioral output leading to sickness phenotypes remains limited. Most experiments have relied on administration of lipopolysaccharide (LPS), pro-inflammatory cytokines and activators, or inhibitors of inflammatory cytokine production to gauge the effects of immune disturbance on animal and human behavior as associated with sickness ^1, 3, 8, 23–26^. Other studies have used transgenic mice to show that CNS-driven overexpression of specific pro-inflammatory cytokines was sufficient to cause structural and behavioral changes mimicking human neurological disorders or sickness^27^. Here, we use an unbiased large-scale genetic approach to examine whether a null mutation in the NOD-like receptor *nlrc3l* causing peripheral macrophage dysregulation and systemic inflammation^28^ can modify homeostatic zebrafish behavioral outcomes. Free and exploratory whole-body movements, coined as “spontaneous locomotion”, represent an innate behavior that does not require learning or prior experience. Because they are arguably the most fundamental and evolutionarily conserved behavioral readout^29–35^, we focused on this mode of behavior. Spontaneous locomotion may also reflect the motivational state and survival instincts of animals to explore, search for food, and respond to environment^35–38^, which are conditions that may be altered by sickness. To this end, we examined daily routine spontaneous swimming continuously over several days in larval zebrafish and assessed the impact of innate immune perturbations.

The larval zebrafish model system offers clear advantages for investigating complex interactions between the immune and nervous systems affecting brain circuits and behavioral outcomes. Due to its optical transparency, small-size, rapid development, and genetic tractability in the intact organism, larval zebrafish is amenable to in-depth in vivo analysis of molecular and cellular processes, allowing mechanisms at the single-cell level to be revealed in the context of the whole animal physiology and behavior. Furthermore, zebrafish genes and immune system share high homology with those of humans^39–41^. It also presents a unique opportunity to study the innate immune system independent of adaptive immune contributions, since adaptive immunity does not become functionally mature until juvenile adult stages^42^.

In this study, we leverage the strengths of the zebrafish to interrogate the ability for peripheral inflammatory macrophages to modulate steady state brain circuits controlling spontaneous locomotion. Zebrafish larvae naturally make short bursts of movement referred to as swim bouts throughout a 24-hour day, albeit movement is minimized in the dark period at night for these diurnal animals^43, 44^, providing an ideal system to monitor spontaneous locomotion. We hypothesized that modulation of neural circuits by the immune system may be one of the intrinsic factors that govern spontaneous behaviors particularly in unprovoked locomotion, which has remained unexplored. Furthermore, we focused on the analysis of a member of the cytoplasmic nucleotide binding oligomerization domain (NOD)-like receptors (NLRs), which are pattern recognition receptors that respond to microbial molecules and danger- and stress-associated molecular patterns^45^. While most NLRs activate pro-inflammatory signaling, some members function as negative regulators of immune response^45^, including *nlrc3l*, a particularly interesting NLR as its deletion causes spontaneous unwarranted systemic inflammation in zebrafish^28^. Microglia, the brain-resident macrophages, are absent^7, 28^, because peripheral primitive macrophages from which they are derived from^46, 47^ are inappropriately activated and diverted from their intended developmental course. Therefore, the study of *nlrc3l* mutants in addition to three other independent microglia-lacking mutants (gene deletions of interferon regulatory factor *irf8*^48^, the xenotropic and polytropic retrovirus receptor *xpr1b*^49^, and myeloid transcription factor *pu.1/spi1b*^50^) enabled a more systematic analysis of the impact of microglia on steady state locomotor circuitry. Taken together, the collective results from this study indicate negligible impact of microglia loss alone, but critical influences of peripheral macrophage-evoked inflammation compounded by microglia deficiency on triggering behavioral changes in *nlrc3l* mutants.

## Results

### A genetic paradigm for assessing impact of inappropriate inflammatory macrophage activation

Previous forward genetic screens have identified loss-of-function mutations in a novel NOD-like receptor, *nlrc3l*, that cause microglia loss due to abnormal macrophage activation in zebrafish^28, 51^. To address possible neuroimmune interactions in motivating locomotor behaviors, we therefore sought to use *nlrc3l^st73^* null mutants as a genetic model to interrogate whether disruptions in macrophages and microglia can impact the locomotor circuitry.

To characterize the immune dysfunction in *nlrc3l* mutants more comprehensively, we conducted an RNA-seq based transcriptome analysis to define differentially expressed genes in the *nlrc3l* mutants compared with their co-housed wild-type and heterozygous siblings at steady state (Fig. 1a). Gene ontology analysis of the 245 genes significantly upregulated in the *nlrc3l* mutants revealed highly significant enrichment of pathways associated with innate immune activation and inflammation, including cytokines (*il1b*, *cxcl8a*), chemokines (*cxcl8a*, *cxcl18b*, *cxcl19*), enzymes (*mmp9*, *mmp13a*, *timp2b*, *irg1*, *irg1l*, *adam8a*, *irak3*, *ncf1*), receptors (*il6r*, *gpr84*, *nlrp16*), and transcription factors (*irf8*, *irf1b*)^52^ known to be elevated during pro-inflammatory conditions (Fig. 1a). *tnfa* was also found to be elevated in *nlrc3l* mutants but not above the threshold of a 1.5 fold increase as previously shown^28^. Conversely, 113 genes were significantly downregulated in *nlrc3l* mutants that largely affect metabolic processes, or notably, are known to be microglia gene markers (*mrc1b*, *lgals3bpb*, *havcr1*)^53^ consistent with microglial cells being absent in *nlrc3l* mutants (Fig. 1a).

**Figure 1.**
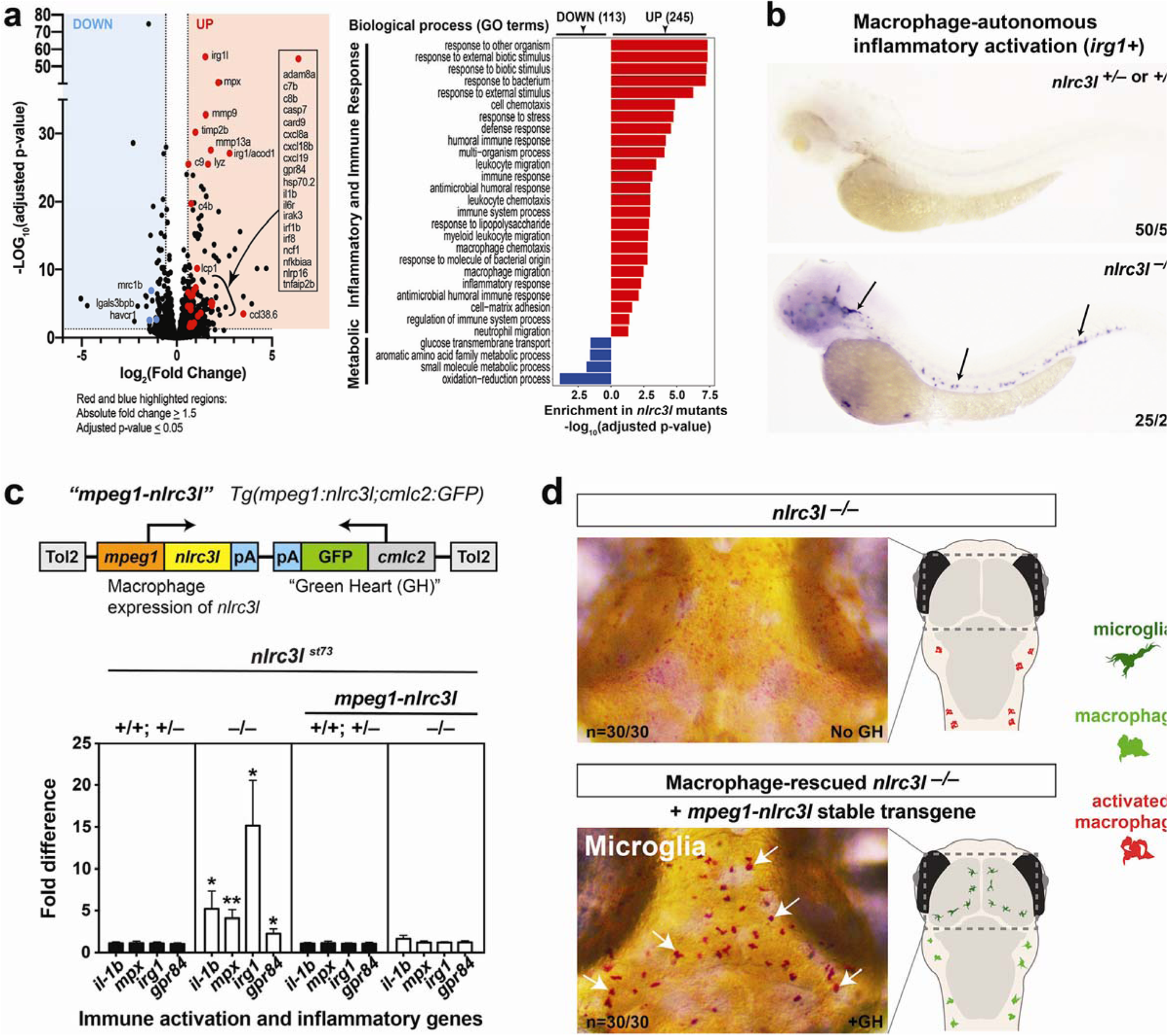
*nlrc3l* knockout causes inappropriate macrophage activation that leads to systemic inflammation and prevents microglia development in zebrafish. (a) RNA-seq analysis comparing *nlrc3l* mutants over heterozygous and wild-type siblings at the 4 dpf larval stage show significant upregulation of inflammatory and immune response genes and downregulation of metabolic and microglia genes in whole larvae. (b) Whole mount in situ hybridization of macrophage activation marker *irg1/acod1* mRNAs in the 2.5 dpf zebrafish larvae show specific and robust induction of *irg1* expression in *nlrc3l* mutants but no expression in wild-type or heterozygous siblings. (c) Top, schematic of the macrophage rescue construct used to generate the stable *mpeg1-nlrc3l* transgenic line to restore wild-type macrophages in *nlrc3l* mutants using the Tol2 transposon system. Bottom, qPCR analysis demonstrates efficacy of the stable macrophage rescue transgene to reverse increased expressions of pro-inflammatory and neutrophil genes in *nlrc3l* mutants, thereby abrogating systemic inflammation Error bars show sem; **, p-value < 0.01; *, p < 0.05. qPCR was conducted using 3 technical replicates and a minimum of 3 biological replicates. (d) Characterization of macrophage-rescued *nlrc3l* mutants demonstrates restoration of microglia in all mutants analyzed carrying the *mpeg1-nlrc3l* transgene (+GH) but not in mutants without the macrophage rescue construct (no GH). Left, neutral red staining marks microglial cells in red (arrows). Right, cartoons of baseline and macrophage-rescued mutants showing status of microglia and peripheral macrophages; dotted box shows region depicted in the neutral red images on the left.

Of particular interest, *acod1/irg1*, aconitate decarboxylase 1 or immune-responsive gene 1, was one of the most highly upregulated genes by RNA-seq analysis in *nlrc3l* mutants. This gene encodes for a conserved mitochondrial enzyme that catalyzes the formation of itaconate, known to be highly elevated in inflammatory macrophages ^54–56^. Similar to mouse, *irg1* is specifically expressed in activated macrophages after bacterial infection, LPS activation, and cancer cell exposure in zebrafish, but not normally in unchallenged wild-type ^54, 57, 58^. By RNA *in situ* hybridization, we found a strong transcriptional induction of *irg1* in macrophages of all *nlrc3l* mutants analyzed, in stark contrast to siblings devoid of any *irg1* expression (Fig. 1b). The spontaneous induction of *irg1* in *nlrc3l* mutants in the absence of infection, injury, or any overt challenge shows that macrophages are indeed inappropriately activated.

Furthermore, dysregulated inflammatory macrophages were implicated to be responsible for the systemic inflammation observed in *nlrc3l* mutants^28^, but direct evidence was not demonstrated. To examine this connection, we created a stable transgenic line to constitutively restore wild-type *nlrc3l* expression in a macrophage-specific manner using regulatory sequences for a macrophage gene *mpeg1* in *nlrc3l* mutants (subsequently referred to as macrophage rescue)(Fig. 1c-d). This macrophage rescue cassette included a cardiac myosin *cmlc2* gene promoter controlled GFP reporter (*cmlc2:GFP*) to enable selection of the transgenic embryos based on heart GFP (GH+) expression. *nlrc3l^st73^* heterozygous intercrosses were used throughout the study to breed homozygous *nlrc3l* mutants and their siblings. Using this conditional macrophage rescue strategy, we show that indeed restoring wild-type macrophages was sufficient to eliminate the significant elevation of pro-inflammatory genes (*il1b, mpx, irg1, gpr84*), including the neutrophil gene marker *mpx*, in *nlrc3l* mutants, thereby reversing systemic inflammation (Fig. 1c). This was associated with a full recovery of microglia in all *nlrc3l* mutants carrying the macrophage rescue transgene (GH+) (Fig. 1d). By contrast, *nlrc3l* mutant siblings without the rescue transgene showed the expected inflammation and loss of microglia phenotypes (Fig. 1c-d).

Taken together, the results from RNA-seq, *acod1/irg1 in situ* expression, and conditional macrophage rescue collectively provide direct evidence that the mutant macrophages are inappropriately activated and responsible for systemic inflammation and microglial absence in *nlrc3l* mutants as previously implicated^28^. These mutants thereby provide an effective genetic paradigm for assessing the impact of inappropriate macrophage-evoked inflammation on other body systems, particularly the brain.

### NOD-like receptor *nlrc3l* mutants capable of normal motility and visuomotor response appear less motivated to move

In the absence of any overt stimuli, locomotion of freely behaving animals represents a major form of spontaneous behavior, possibly reflecting some degree of intrinsic motivation for exploration and physical activity^59–61^. To examine whether inappropriately activated macrophages could disrupt spontaneous behaviors in larval zebrafish, we analyzed *nlrc3l* mutants using a large-scale automated tracking system equipped with a high-speed infrared CCD camera to simultaneously monitor freely swimming individually-housed zebrafish larvae at 5 days post-fertilization (dpf) under normal day-night cycles in 96-well (Fig. 2a and Supplementary Movie 1) and 24-well platforms (about 5× larger volume than the 96-well) (Fig. 2a and Supplementary Movie 2). The larger housings enabled analysis of not only motility but also spontaneous swimming patterns (Fig. 2a) that may be exploratory in nature. Animals were tracked simultaneously in their chambers for three continuous days over the normal 14-hour light and 10-hour dark cycle with all genotypes blinded (Fig. 2a).

**Figure 2.**
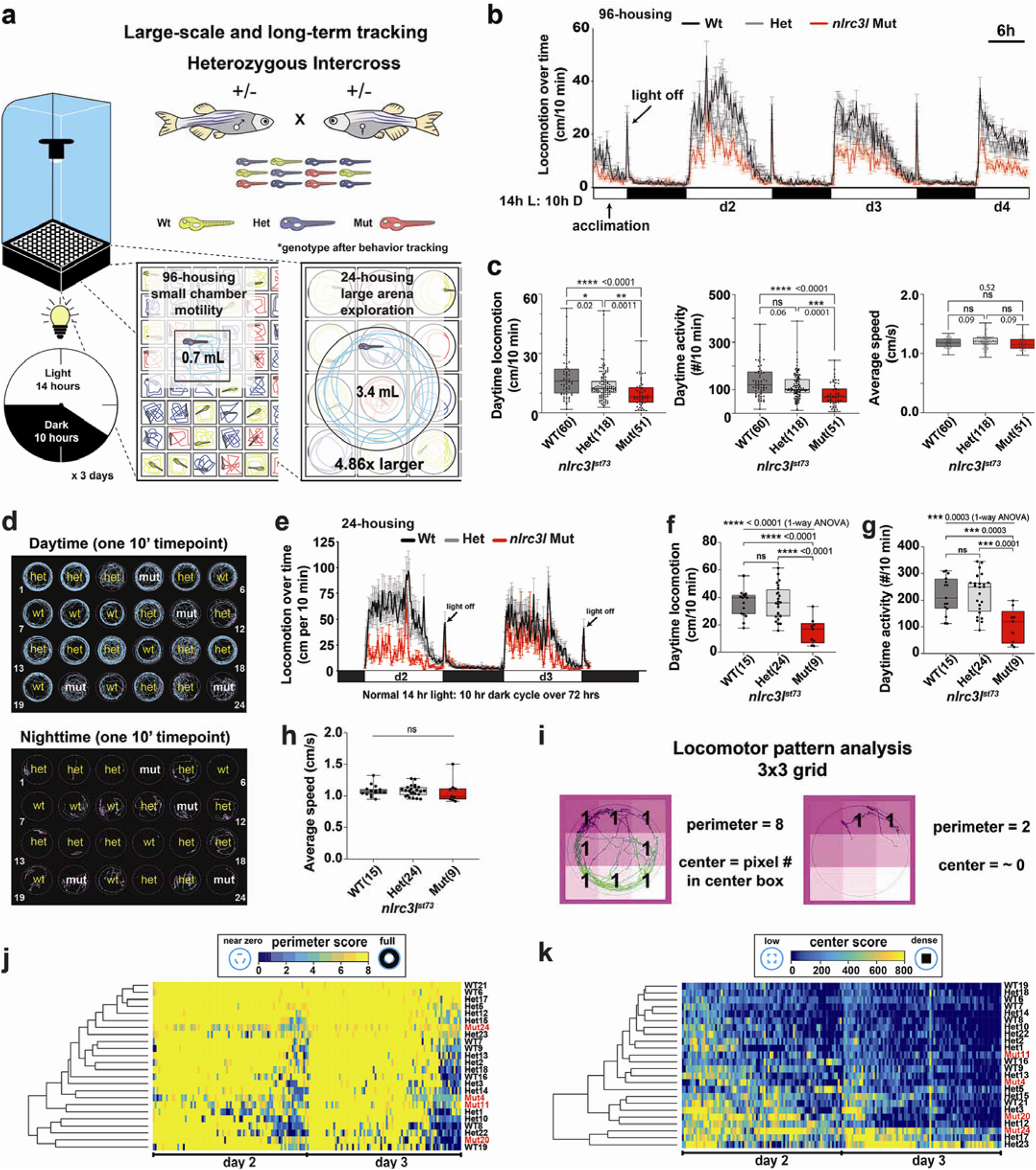
Long-term behavior tracking reveals a significant reduction in daytime locomotion in *nlrc3l* mutants due to reduced number of swimming bouts in both 96-well and 24-well chambers. (a) Schematic of the automated behavior tracking system for 96-well and 24-well chamber setups. Mutants are derived from heterozygous intercrosses and all animals are randomized and individually placed in each well without prior knowledge of their genotypes. See recording of animals in 96-well (Supplementary Movie 1) and in 24-well (Supplementary Movie 2) housings. (b) Representative time plot of locomotor tracking in 96-well chamber. 14-hour light cycle and 10-hour dark cycle (14h L: 10h D) was applied. Light turns on at 7 am and turns off at 9 pm each day, and the “light off” event triggers the expected hyper visuomotor response (sudden rapid increase in swimming activity, arrow). *nlrc3l* mutants (red graph) consistently show a significant average decrease in spontaneous swimming. (c) Calculations of different daytime locomotor metrics from three independent behavior tracking sessions. One of these sessions is represented in time plot in b. (d) Representative swimming traces of a day and a night timepoint from 24-well tracking. Genotypes were determined after completion of experiment and added back to the traces. Traces show typical inter-individual variation not readily associated with a genotype. Activity is highly suppressed at night. Teal traces mark large movements (≥0.5 cm/s) and magenta traces mark small movement (< 0.5 cm/s). (e-h) Behavior tracking using the large-arena 24-well platform and associated quantifications from two independent behavior tracking sessions. See associated Supplementary Movie 3. (i) Diagram illustrating locomotor pattern analysis based on a 3×3 grid system. Traces of swimming trajectories were recorded at every 10-minute interval over the entire 72-hour tracking. Traces corresponding to d2 and d3 timepoints were scored. See Materials and Methods for details on the calculations. (j-k) Heatmaps combined with hierarchical clustering of perimeter and center scores do not show any segregation of fish based on genotype, indicating similar patterns of locomotion between *nlrc3l* mutants and their siblings. (c, e-h) Number in parenthesis is *n*, number of individual animals analyzed. Scatter box-and-whisker plots show minimum and maximum; each data point represents an individual animal. Daytime locomotion is measured by the total swimming distance per 10-minute interval. Daytime activity is number of bouts of swimming detected per 10-minute interval. All metrics were averaged over d2 and d3 timepoints per fish. d2, 14-h light period on day 2; d3, 14-h light period on day 3; d4, partial light period on day 4. Scale bar, 6h (hours). ns, not significant; *, p<0.05; **, p<0.01; ***, p<0.001; ****, p<0.0001; all p-values are FDR-adjusted. Statistical significance was calculated using multiple comparisons test after one-way ANOVA test, or the Brown-Forsythe and Welch ANOVA test for groups with unequal standard deviations.

Using this long-term tracking, we found a striking reduction in spontaneous daytime swimming in *nlrc3l* mutants in both 96-well and 24-well housings. These mutants exhibited the expected high activity during the day and periods of resting at night of diurnal animals (Fig. 2b). In the 96-well format, we found significantly large reductions of 35-50% in median daytime distance travelled and 30-40% in median frequency of swimming activity in *nlrc3l* mutants, while no difference in average swimming speed compared with heterozygous and wild-type siblings (Fig. 2b-c). When comparing large numbers between heterozygous and wild-type control animals (each n>50), small significant locomotor differences could be detected for distance travelled (Fig. 2c), but not apparent in smaller sample sizes (Fig. 2f), suggesting the difference, if any, may be trivial. Large arena 24-well analysis also showed reduced spontaneous daytime swimming in *nlrc3l* mutants (Fig. 2d-h). Both housing platforms indicate that a lower number of swimming bouts (daytime activity) accounted for the reduced locomotion as the average speed of swimming was unchanged.

While *nlrc3l* mutants are not visibly different from wild-type either by gross physical or behavioral traits (no obvious defect in skeletal muscles, response to touch, or general motility) or by traces of movement at any given timepoint, we sought to analyze the quality of their swimming in more detail. Swimming traces dynamically varied within and between individuals (Fig. 2d and Supplementary Movie 3), so analyzing a large number of animals over a long time period was imperative to acquire reliable trends. We developed a quantitative method to characterize swimming patterns in the large 24-well arenas based on image segmentation of swimming traces from 14-hour light periods on days 2 and 3 (d2 and d3) using a 3 x 3 grid system (Fig. 2i). A heatmap of the perimeter and center scores of individuals tracked over time conveyed the inter- and intra-individual variation in their swimming pattern (Fig. 2j-k and Supplementary Fig. 1). In general, locomotor activity was high at the beginning of the day and declined toward the end of the day (Fig. 2j-k). By unsupervised hierarchical clustering of perimeter (Fig. 2j) and center (Fig. 2k) scores, we found *nlrc3l* mutants to be sorted and grouped together with control siblings, thereby indicating similar swimming patterns for all genotypes. Based on the fraction of the daytime spent on full or no perimeter laps, and high or near zero center locomotion, *nlrc3l* mutants were similar to control siblings except for a significant 30% reduction in swimming around the full perimeter (Supplementary Fig. 1). We found anti-correlated trends between frequency of full perimeter swimming and low center activity in all genotypes, indicating that more perimeter swimming coincided with more center swimming (Supplementary Fig. 1). Taken together, behavioral tracking in small and large housings and the swimming pattern analysis indicate that *nlrc3l* mutants are capable of normal swimming, but conduct less perimeter swimming than their wild-type counterparts.

To determine whether impairment in visuomotor coordination may contribute to the *nlrc3l* mutant behavior, we tested these fish on a rapid light-to-dark switch assay known to activate a stereotypical hyper-swimming response^62–64^ (Supplementary Fig. 2). We found that the *nlrc3l* mutants responded to a light-to-dark switch with an equivalent rapid increase in swimming as their wild-type counterparts (Supplementary Fig. 2), indicating normal motility and no visuomotor impairment. Interestingly, when we subjected the same fish cohort to long-term 72-hour tracking after the visuomotor test, a reduction in daytime locomotion resumed in *nlrc3l* mutants (Supplementary Fig. 2). Collectively, our data demonstrate that *nlrc3l* mutants are capable of normal motility and visuomotor coordination, but appear less motivated to swim when unprovoked. These results implicate a disruption of the upstream supraspinal circuits in *nlrc3l* mutants underlying the brain control of locomotion rather than the effector spinal circuits that execute locomotion.

### Aberrant inflammatory macrophages causing systemic inflammation lead to reduced daytime locomotion

In light of the known macrophage defects in *nlrc3l* mutants, we asked whether the inappropriately activated peripheral macrophages were responsible for the reduced locomotor activities. We turned to using the stable macrophage rescue line expressing *mpeg1-nlrc3l* as described above (Fig. 1c) to determine whether restoring wild-type macrophages in *nlrc3l* mutants was sufficient to re-establish normal levels of spontaneous swimming. A large-scale 96-well platform was used to test 48 individuals with the macrophage rescue transgene (GH+) and the remaining 48 wells were used for siblings without the transgene (non-rescued, no GH) as negative controls (Fig. 3a). All fish were derived from a heterozygous intercross for the *nlrc3l^st73^* mutation with one parent carrying the macrophage rescue transgene. We analyzed the locomotor behaviors in the normal light-dark cycles over a continuous 72-hour period and found the expected reduction in daytime swimming in control *nlrc3l* mutants without the rescue construct (no GH)(Fig. 3b-c). By contrast, the macrophage-rescued *nlrc3l* mutants showed a significant recovery of daytime locomotion akin to wild-type and heterozygous animals (Fig. 3b-c). We did not detect any difference in locomotor behaviors in wild-type or heterozygous individuals due to expression of the macrophage-rescue *mpeg1:nlrc3l* transgene (Fig. 3b-c). Furthermore, we found that the macrophage-rescued *nlrc3l* mutants also significantly recovered exploratory behaviors in the large-arena 24-well tracking (Supplementary Fig. 3). In the large-arena environment, the rescue transgenic expression also did not alter swimming levels of control wild-type and heterozygous individuals (average distance (cm) travelled per 10 min ± standard deviation was 42.0 ± 11.5 for GH+ WT and 46.3±18.8 cm/10 min for GH+ Het which overlapped in range as non-transgenic WT at 35.6 ± 9.9 and non-transgenic Het at 36.4 ±12.3) (Supplementary Fig. 3 and Fig. 2f). By contrast, macrophage-rescued GH+ mutants restored wild-type levels of swimming in the large arena at 36.7 ± 8.9 cm per 10 min compared with their significantly reduced baseline swimming at 15.5 ± 9.6 cm per 10 min in the non-transgenic mutants (Supplementary Fig. 3 and Fig. 2f). Taken together, these results show that restoring wild-type macrophages in *nlrc3l* mutants was not only sufficient to recover microglia and reverse inflammation (Fig. 1c-d), but also significantly improves levels of daytime swimming in *nlrc3l* mutants (Fig. 3 and Supplementary Fig. 3). The data indicates that the inappropriately activated peripheral macrophages were responsible for the reduced locomotor behaviors in *nlrc3l* mutants, underscoring a possible innate immune modulation of the brain (or supraspinal) circuits that regulate spontaneous locomotion.

**Figure 3.**
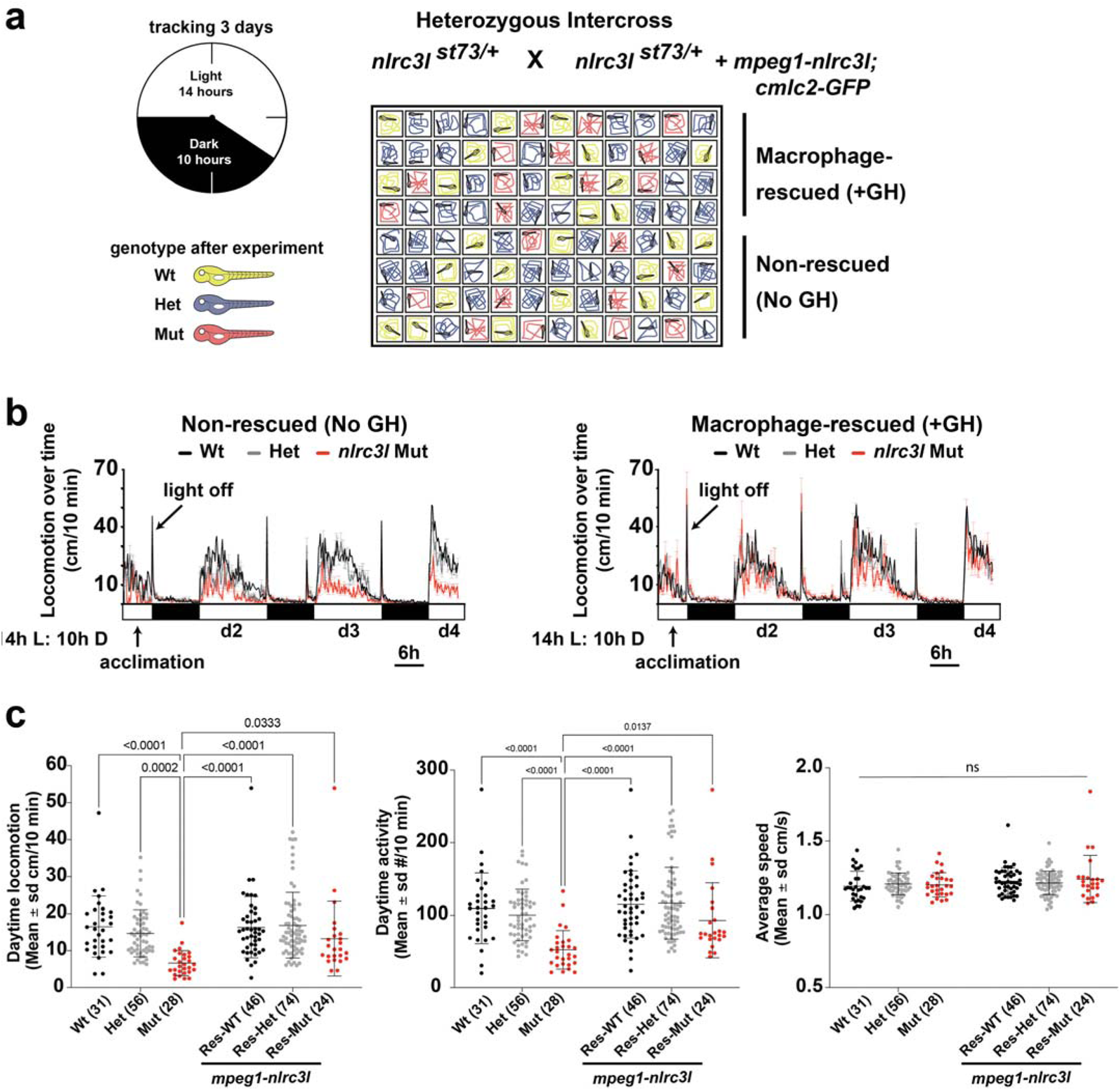
Macrophage-specific rescue of *nlrc3l* mutants is sufficient to reverse the deficient locomotor phenotype. (a) Schematic of experimental setup to analyze locomotor behaviors of macrophage-rescued (+GH) and non-transgenic non-rescued (No GH) sibling zebrafish side by side. (b) Time plots showing the efficacy of the stable *mpeg1-nlrc3l* transgene to largely restore daytime locomotion in *nlrc3l* mutants from their reduced baseline level (shown in the non-rescued plot). The rescue transgene had no apparent effect on the heterozygous and wild-type siblings; no difference in locomotor behavior was found between siblings with and without the rescue transgene. (c) Scatter box-and-whisker plots showing no significant difference in daytime locomotion or activity in macrophage-rescued *nlrc3l* mutants compared with baseline wild-type siblings, while baseline *nlrc3l* mutants showed a significant reduction. *n*, number of animals analyzed shown in parenthesis below each scatter plot. Three independent experiments were conducted to compare larvae derived from a *nlrc3l* heterozygous intercross with and without the macrophage rescue transgene on the same plate. Statistical significance was determined by two-way ANOVA test followed by multiple comparison tests. FDR-adjusted p-values are reported; ns, not significant as defined by p-values > 0.05. Pairwise brackets not shown are ns, not significant. Res, individuals carrying the rescue transgene.

### Absence of microglia cannot solely account for the reduced daytime locomotion

Because microglia are implicated in synaptic pruning and modulation of neuronal circuit function^17, 65, 66^, we investigated whether microglia loss in *nlrc3l* mutants may be another major cause of their locomotor deficit, in addition to the inappropriate macrophage activation. We found microglia widely distributed throughout the brain in wild-type zebrafish at 5 dpf at the time of our behavioral analyses, including in the deeper regions of the telencephalon and diencephalon that are known to control locomotor circuits^67–69^, besides their well-known tectal localization^70^ (Fig. 4a and Supplementary Movie 4). In addition to constant interactions with tectal neurons, microglia intimately intermingled with neurons in the ventral regions of the telencephalon and diencephalon (Fig. 4b and Supplementary Movie 5). The broad distribution of microglia is consistent with the possibility that these cells can impact development or function of various brain circuits.

**Figure 4.**
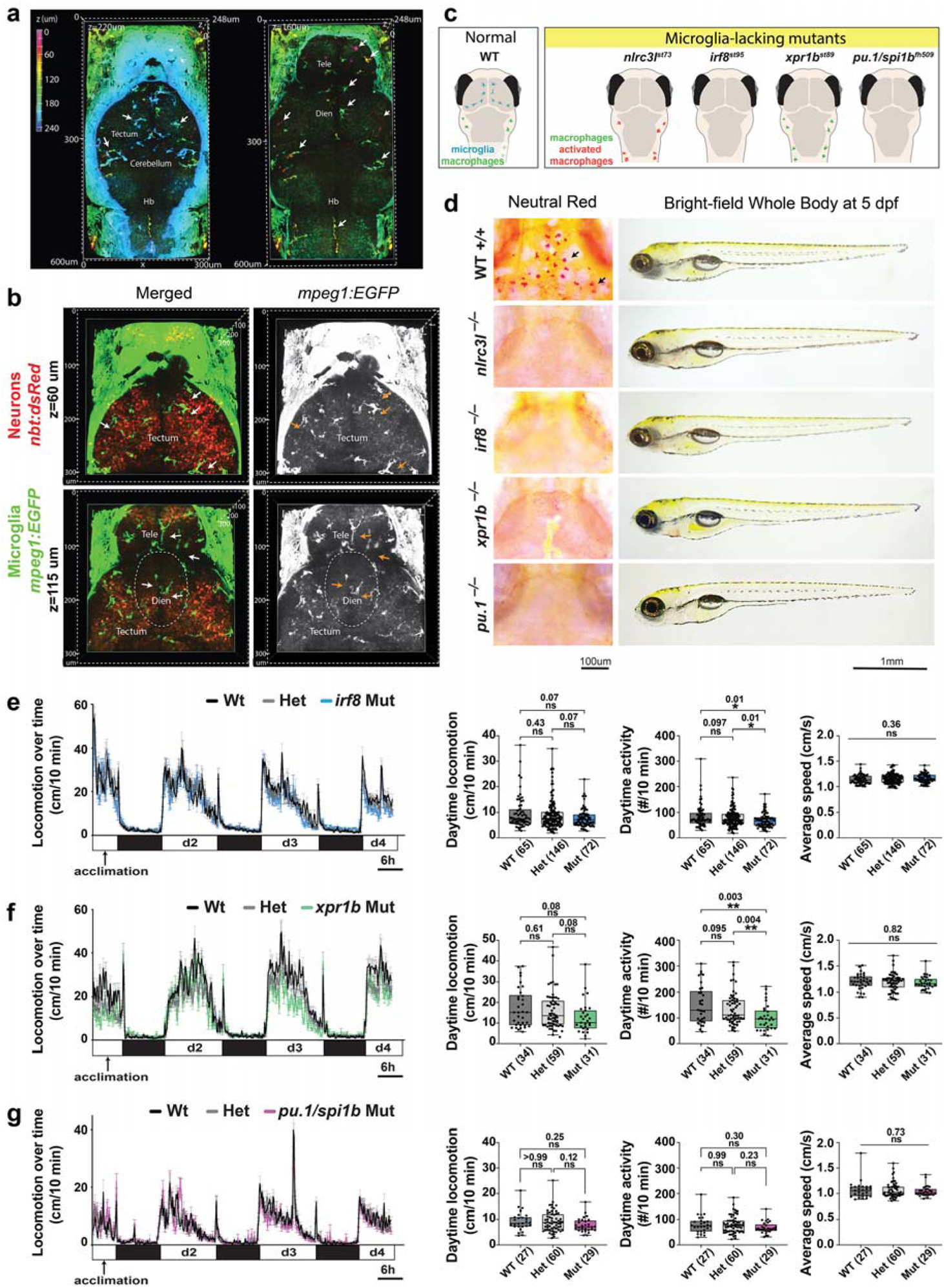
Microglia intermingle with neurons throughout the healthy larval zebrafish brain, but have minimal to no impact on spontaneous locomotion. (a) Volumetric views of wild-type 5 dpf larval brain showing microglia (arrows) using the *mpeg1:GFP* transgene as cells depth-coded from most ventral in magenta to most dorsal in cyan. See corresponding z-stack data in Supplementary Movie 4. Left, dorsal surface at z=220 um shows the tectum, cerebellum and hindbrain (hb). Right, more ventral view at z=160 um shows the telencephalon (tele) and diencephalon (dien). (b) Volumetric views of a z-stack from two axial levels (z=60 um and 115 um) of a wild-type double transgenic zebrafish at 5 dpf showing microglia (GFP+) intimately intermingled with neurons (dsRed+) throughout the larval brain. See corresponding z-stack data in Supplementary Movie 5. (c) Diagram depicting microglia and peripheral macrophage status in wild-type and four distinct microglia-lacking mutants in larval zebrafish. (d) Characterization of the four mutants (*nlrc3l*^−/−^, *irf8*^−/−^, *xpr1b*^−/−^ and *pu.1/spi1b*^−/−^) by neutral red staining shows an absence of microglia, and by bright-field transmitted microscopy shows normal gross morphology with large swim bladders indistinguishable from wild-type. Locomotor behavior tracking in the 96-well platform analyzing *irf8* mutants (e), *xpr1b* mutants (f), and *pu.1/spi1b* mutants (g). One-way ANOVA tests followed by multiple comparisons were used to determine statistical significance; individual p-values shown; locomotion from d2-d4 was analyzed; ns, not significant. See also associated Supplementary Movies 4 and 5.

Since uncoupling microglia loss from inflammatory processes in *nlrc3l* mutants is unattainable because both are consequences of the early inappropriately activated macrophages, we investigated whether an absence of microglia alone is sufficient to phenocopy *nlrc3l* mutant locomotor behavior. To this end, we extended our behavioral analysis to three other microglia-lacking mutants (*irf8*^−/−^, *xpr1b*^−/−^, and *pu.1/spi1b*^−/−^), which do not have the inflammation phenotypes^48–50, 71^(Fig. 4c). All three mutants lack microglia due to developmental disruptions distinct from *nlrc3l* mutants (Fig. 4c-d); both *irf8* and *pu.1/spi1b* are conserved transcription factors required for myeloid and macrophage cell development^48^ whereas *xpr1b* is a phosphate exporter required for migration and differentiation of microglial precursors^49^. In common, all four microglia-lacking mutants including *nlrc3l^st73^* do not exhibit any gross morphological defects with normally sized swim bladders, and are anatomically indistinguishable from wild-type fish (Fig. 4d). Time plot analysis and calculations of daytime locomotion surprisingly indicate that *irf8* (Fig. 4e), *xpr1b* (Fig. 4f), and *pu.1* (Fig. 4g) mutants were largely normal relative to control siblings, showing no significant difference in locomotion based on distance of movements, although a rather modest decrease in daytime activity count (average median reduction of 14%) for *irf8* and *xpr1b* mutants (Fig. 4e-f). This is a relatively small change in comparison to the large average median reduction of 36.5% in *nlrc3l* mutant daytime activity (Fig. 2c), indicating that an absence of microglia alone could not explain the locomotor deficit in *nlrc3l* mutants.

### Pro-inflammatory conditions are causal to disrupted daytime locomotion

Given the implications of pro-inflammatory factors on causing reduced locomotion during sickness^3, 5^ and inflammation as possibly a main driver of the locomotor deficiency in *nlrc3l* mutants, we tested whether reversing systemic inflammation by administering anti-inflammatory drugs could significantly restore normal behaviors in *nlrc3l* mutants. We examined three small-molecule drugs (17-DMAG, Bay 11-7082, and dexamethasone) that are water-soluble and compatible for administering in the fish water (Supplementary Fig. 4). They have been shown to curb inflammation through different mechanisms by attenuating NF-kB mediated transcription or activating glucocorticoid functions in zebrafish and other systems. 17-DMAG is a water-soluble geldanamycin analog that can inhibit the heat-shock protein Hsp90 and cause degradation of target proteins, including the NF-kB protein complex^72, 73^; Bay 11-7082 can inhibit E2 ubiquitin (Ub) conjugating enzymes and prevent degradation of IkB-alpha, an NF-kB inhibitor^74^; and dexamethasone is an agonist of the glucocorticoid receptor (GR) that activates a negative feedback mechanism to reduce inflammation^75^. We used the 96-well platform to enable a large sample size to test the different drugs at concentrations previously characterized to reduce inflammatory processes^71^. Each drug administered was tested side-by-side on the same plate with a control (no treatment) condition on fish derived from the same clutch of a *nlrc3l^st73^* heterozygous intercross (Supplementary Fig. 4). We found that treatment with dexamethasone, but not 17-DMAG or Bay 11-7082, was effective in significantly restoring daytime locomotion in *nlrc3l* mutants to levels similar to untreated control siblings (Supplementary Fig. 4). The difference in the effect of the three drugs suggest that the mutants may be deficient in glucocorticoid regulation (which dexamethasone activates) rather than excessive NF-kB activation (which 17-DMAG and Bay-11-7082 inhibit). Interestingly, because dexamethasone appears to increase daytime spontaneous locomotion in control siblings consistent with another previously published study^76^, dexamethasone may have a stimulatory effect in addition to suppressing inflammation. It is, however, important to point out that dexamethasone increased locomotion in mutants more so than in controls as shown by the overlapping range of locomotor activity after dexamethasone treatment, instead of a proportional increase. Furthermore, additional means of suppressing inflammation by restoring macrophage *nlrc3l* function also significantly restored locomotor activity in *nlrc3l* mutants. Taken together, these results indicate that administering dexamethasone, a synthetic glucocorticoid, without restoring microglia or reversing macrophage inflammation was effective to reverse a significant level of the *nlrc3l* mutant locomotor deficit, lending support for pro-inflammatory signaling being a predominant factor in disrupting the corresponding locomotor circuits.

### Reduced locomotion is associated with dorsal thalamic calcium imbalance

To define brain regions that may be disrupted in *nlrc3l* mutants, we used genetically encoded calcium indicator GCaMP6 fast variant (GCaMP6f) to assess four brain regions known *in vivo* to either contain circuits controlling locomotion in vertebrates (cerebellum, diencephalon (dorsal thalamus), and dorsal telencephalon (pallium))^34, 77, 78^ or a high density of microglia (tectum)^70, 79^ (Fig. 5a). Due to the implicated link between microglia and neuronal development^17, 65, 66^, we hypothesized that brain regions in or near the proliferative zone or ventricular surface may be more susceptible to immune interference or modulation. We therefore focused our analysis on a subset of newly differentiated neurons using the basic helix-loop-helix (bHLH) transcription factor *neurod* regulatory sequence^80–82^ to drive GCaMP6f in these neurons for *in vivo* calcium imaging. We assessed spontaneous neural activity in these regions to determine possible differences underlying the reduced locomotion in *nlrc3l* mutants compared with their control siblings using resonant scanning microscopy at 6 dpf under normal daytime conditions. Interestingly, we found no significant difference in number of calcium transients in the telencephalon, diencephalon, or cerebellum, but a significant increase in tectal neuronal activity in *nlrc3l* mutants compared with control siblings (Fig. 5b-f). To determine whether this increase was due to impaired macrophages in the *nlrc3l* mutants, we asked whether restoring normal macrophages in the *nlrc3l* mutants could reverse this tectal activity increase (Fig. 5b-f). Using the stable macrophage rescue line as described in Fig. 1c, we found indeed this was the case, while neuronal activity in the other brain regions had no apparent changes (Fig. 5b-f).

**Figure 5.**
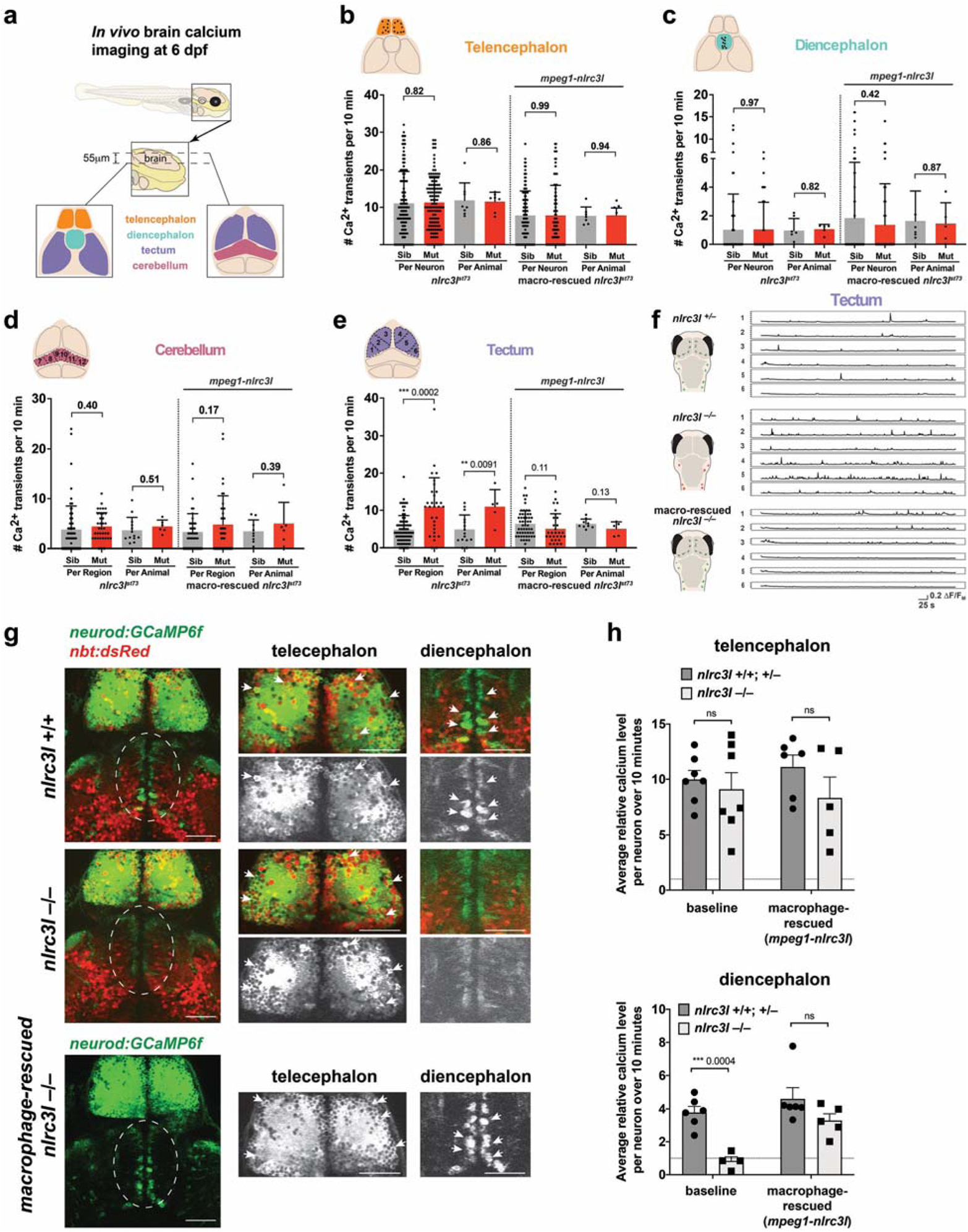
*In vivo* imaging of spontaneous brain neuronal activity reveals increased tectal calcium transients while decreased diencephalic calcium in *nlrc3l* mutants. (a) Schematic showing the brain regions captured by calcium imaging. Two z-planes in the 6 dpf larval brain were imaged using the genetically encoded calcium indicator *neurod:GCaMP6f*. (b-e) Scatter bar charts show the number of spontaneous calcium transients in the corresponding brain regions as illustrated for baseline and after macrophage-rescue. Cerebellum and tectum were each divided into 6 regions for analysis as depicted in each cartoon. As indicated, calcium transients were calculated on an individual neuron or region basis, or per animal basis using the average calcium transient number of the neurons or region of analysis. The only significant difference found in *nlrc3l* mutants compared with control siblings was an increase in tectal neuronal calcium transients, which was reversed after macrophage rescue. (f) Traces from representative individuals of each genotype showing calcium activity *(GCaMP6f+)* from each of the six tectal regions (1-6) analyzed. Plots show an apparent increase in spiking events in the *nlrc3l* mutant at baseline. (g) Relative intracellular calcium levels in the telencephalon and diencephalon (dotted region) at baseline were determined. *nlrc3l* mutants had consistently lower intracellular calcium in diencephalic neurons corresponding to the dorsal thalamus compared with control siblings, but this was reversible by the macrophage rescue. Arrows point to neurons with robust somatic calcium levels (high *GCaMP6f+)*. (h) Bar charts show relative intracellular calcium levels in the telencephalon and the diencephalon as an average over the neurons analyzed for each respective region. Each data point represents an individual animal. No difference was determined in intracellular calcium level in the telencephalic neurons based on genotype, but diencephalic neurons were found to have significantly lower intracellular calcium in *nlrc3l* mutants compared with control siblings. ns, not significant. Two-tailed t-test was used to determine statistical significance. See also associated Supplementary Fig. 8 and Supplementary Movies 6 and 7.

Since the macrophage rescue strategy cannot distinguish recovery of microglia from reversal of systemic inflammation, because both are direct consequences of the mutant macrophages, we next asked whether a sole microglia loss using the microglia-lacking *irf8* mutant^48^ could account for the increased tectal neuronal activity. To address this, we conducted calcium imaging of tectal neurons in the *irf8* mutants, and found an increase in tectal calcium transients compared with sibling controls at both the animal level (p=0.03) and individual neuron level (p=0.06) (Supplementary Fig. 5). These results suggest that microglia loss alone could be responsible for the increase in tectal activity. Consistent with these results, microglia have been shown *in vivo* to negatively regulate neuronal activity in the zebrafish tectum^83^ and mouse dorsal striatum^84^. Since a strongly reduced locomotor behavior was not found in *irf8* mutants, we reason that the common increase in tectal activity in *nlrc3l* mutants and *irf8* mutants was likely not responsible for the *nlrc3l* mutant behavior, but merely reflecting a consequence of their mutual microglia loss.

While the frequency of calcium spikes was not changed, we found a robust decrease in baseline intracellular GCaMP6f fluorescence level in the diencephalic ventricular neurons corresponding to the dorsal thalamus that stood out in the *nlrc3l* mutant brains compared with the siblings (Fig. 5g-h). We quantified the relative change in total cell fluorescence for a random selection of individual neurons in either the telencephalon (as a control) or the diencephalon over a 10-minute period of in vivo calcium imaging (Fig. 5g-h). Our analysis shows that *nlrc3l* mutants had on average ∼4-fold decrease in diencephalic ventricular calcium level compared with their sibling controls, while there was no difference in calcium levels in the telencephalic neurons (Fig. 5g-h). To determine whether this decrease was linked to the mutant macrophages in *nlrc3l* mutants, we used the macrophage-rescue construct and found that the calcium level decrease was reversed after recovery of normal macrophages (Fig. 5g-h), attributing the neuronal calcium phenotype to the mutant macrophages. We then also analyzed whether another microglia-lacking mutant (*irf8^−/−^)* had reduced diencephalic neuronal calcium but did not find a significant decrease, although a marked reduction can be quantified (Supplementary Fig. 6). Taken together, the significantly large decrease in dorsal thalamic intracellular calcium was unique to the *nlrc3l* mutants, raising the possibility that this neuronal disruption was linked to their altered behavior. This significant variation in baseline intracellular calcium level may reflect changes to intra-neuronal calcium signaling and homeostasis, thereby altering excitability of the neurons^85^.

### Disruption of diencephalic dopaminergic TH levels is consistent with reduced locomotion

The severely reduced baseline calcium level of diencephalic neurons in *nlrc3l* mutants points to a possible disruption of the diencephalic circuit as a cause for the behavioral deficit. In support of this, dopaminergic neurons in the diencephalon are known to regulate brain and spinal cord motor circuits in zebrafish and mammals^86, 87^. To assess whether dopaminergic neurotransmission in the diencephalon may be altered in *nlrc3l* mutants, we analyzed possible changes to expression of dopaminergic system genes, namely the two tyrosine hydroxylase TH genes (*th* and *th2*) encoding the rate-limiting enzyme for dopamine synthesis and the dopamine transporter *dat/slc6a3* essential for reuptake of dopamine to terminate neurotransmission^88^. Interestingly, using qPCR, we found that *nlrc3l* mutants had significantly reduced levels of *th* but not *th2* or *dat* (Fig. 6a), raising the possibility that these mutants may have a defect in dopamine synthesis or reduction in number of dopaminergic neurons. Since expression of *th2* is restricted to a small subset of dopaminergic neurons whereas dopaminergic neurons broadly express *th*^89^, the specific reduction in *th* corroborates the possible alteration in the dopaminergic diencephalic clusters that are known to regulate locomotion^69, 90^ in *nlrc3l* mutants. Furthermore, the *dat* gene expression pattern largely overlaps with *th* but was not reduced in *nlrc3l* mutants suggesting that the number of dopaminergic neurons or recycling of dopamine may not be altered.

**Figure 6.**
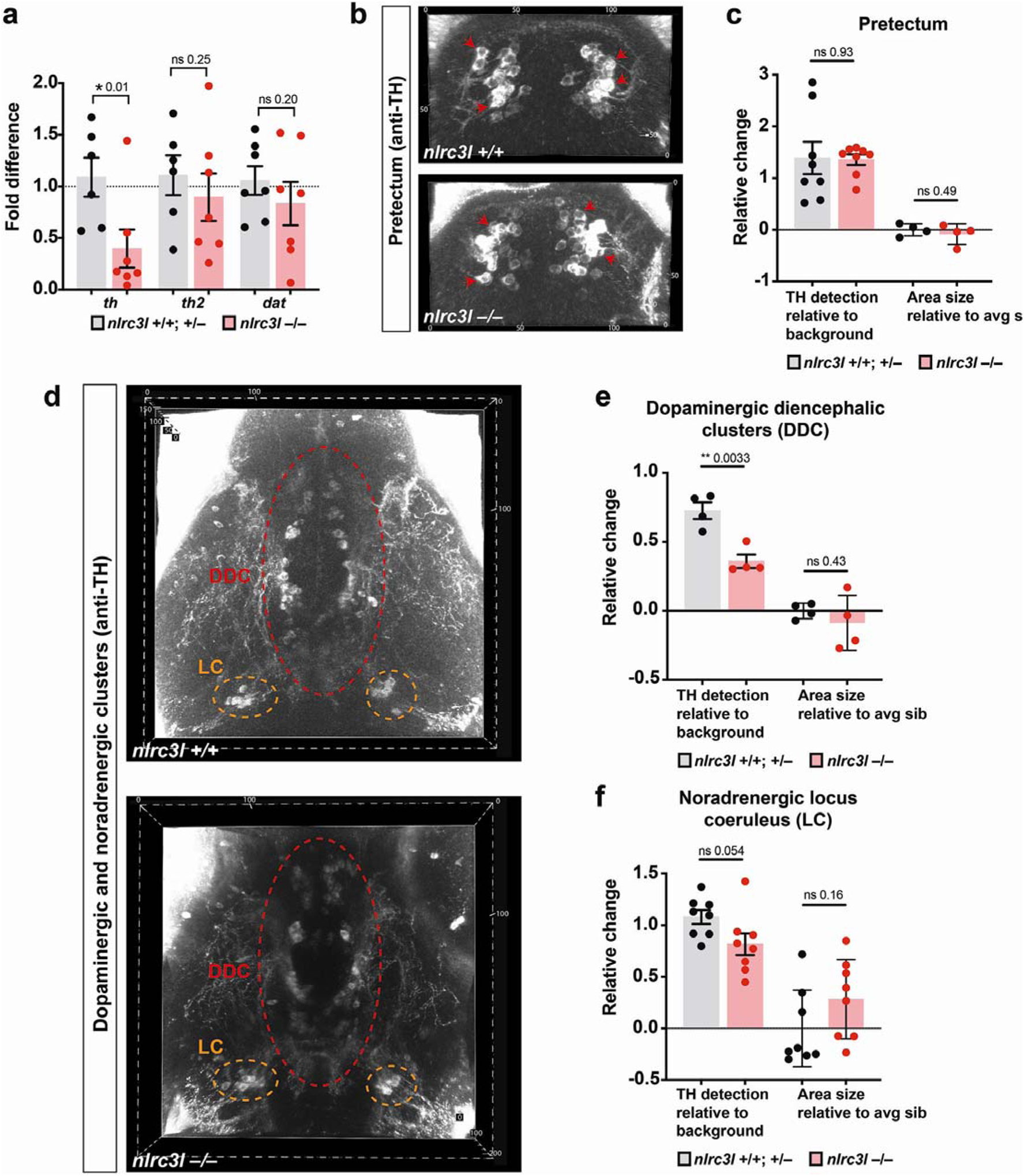
Protein expression of the dopamine synthesis enzyme, tyrosine hydroxylase (TH), is strongly reduced in the dopaminergic diencephalic clusters (DDC) in *nlrc3l* mutants. (a) Bar chart conveys qPCR analysis of dopaminergic system genes (*th*, *th2* and *dat*), indicating a significant decrease in *th* mRNA expression in *nlrc3l* mutants. (b) Anti-TH immunostaining characterization of the pretectal dopaminergic neurons (red arrows) at 6 dpf shows no apparent difference between *nlrc3l* mutant and control wild-type. (c) Quantification of the relative TH protein level and relative area of TH+ staining. (d-f) Characterization of the ventral larval brain at 6 dpf shows a large significant reduction in TH protein level in the posterior tuberculum of the DDC, and a modest and nearly significant decrease in the noradrenergic locus coeruleus (LC). Right, quantifications of the relative TH protein level and relative area of TH+ staining in the DDC and LC. Two-tailed t-test was used to determine statistical significance. Error bars show sem for relative TH staining and sd for relative area.

To further distinguish whether *th* downregulation signifies a decrease in number of dopaminergic neurons or dopamine synthesis, we turned to investigate the TH protein level using a commercially available antibody specific to TH that does not bind TH2^89, 91^. Using confocal brain imaging and immunofluorescence for the TH protein, we quantified the relative TH protein level and TH-expressing neuronal cluster size in three prominent brain regions (Fig. 6b-f). Co-localization analysis of TH protein expression with the dorsal thalamic neurons (as labeled by *neurod:GFP*+) showed that the *neurod*+ neurons imaged for calcium dynamics are not the same as the TH+ neurons, but may directly interact (Supplementary Fig. 7). To assess activity of the TH+ neurons in the diencephalon, we turned to using a pan-neuronal calcium indicator *elavl3:GCaMP6s* for in vivo imaging at two focal planes, through the dorsal thalamus and the presumptive DDC (Supplementary Fig. 8 and Supplementary Movies 6-7). Consistent with *neurod+* calcium imaging results, the basal intracellular calcium levels in both areas of the diencephalon (dorsal thalamus and the DDC) were strikingly reduced in *nlrc3l* mutants compared with control siblings, while no difference was found in the telencephalon nor in number of calcium transients with the exception of a possible small increase in dorsal thalamic calcium activity (Supplementary Fig. 8). We then focused on two regions that consist of dopaminergic neurons for TH expression analysis: pretectum and dopaminergic diencephalic clusters (DDC), with the latter containing dopaminergic diencephalospinal neurons that directly project to the spinal motor circuits^69^, and the locus coeruleus (LC) which contains TH+ noradrenergic neurons as a comparison group. In *nlrc3l* mutants, the DDC, which has descending projections to the motor circuits, showed a significant reduction in TH protein level alongside a modest and nearly significant TH reduction in the LC (Fig. 6e-f), but there was no change in the other brain regions analyzed. The sizes of the neuronal clusters in all regions of analysis were also not altered, suggesting that the defect in *nlrc3l* mutants was likely in neural function rather than structure. These results in combination with aberrant calcium levels in the diencephalic region suggest that *nlrc3l* mutants may have disrupted diencephalic functions that cause reduced spontaneous locomotion.

Since the RNA and protein levels of TH were reduced but the sizes of the neuronal clusters did not significantly change, dopamine synthesis but not neural structure may be affected in *nlrc3l* mutants. Whole mount brain RNA in situ analysis further affirms a grossly normal brain structure in *nlrc3l* mutants by probing a diverse set of neuronal genes: nuclear receptor *nr4a2b/nurr1* marking a subset of dopaminergic neurons^92^, serotonin transporter *serta/slc6a4a* showing serotoninergic neurons^93^, dopamine beta-hydroxylase *dbh* labeling noradrenergic neurons^94^, vesicular-glutamate transporter *vglut2a/slc17a6b* showing excitatory glutamatergic neurons^95^, and glutamate decarboxylase *gad1b* marking inhibitory GABA neurons^95^ (Supplementary Fig. 9). While the broad expression of *vglut2a* and *gad1b* indicate gross overall similarity between *nlrc3l* mutants and control siblings, we cannot exclude the possibility that detailed differences in the connection or pattern of the excitatory and inhibitory neurons exist. Taken together, given the locomotor deficit in *nlrc3l* mutants being linked to systemic inflammation compounded by an absence of microglia, these results implicate a role for the innate immune system in modulating the diencephalic dopaminergic system that regulates locomotor behaviors.

### Abnormal infiltration of neutrophils in the diencephalic brain disrupts locomotor control

How mutant macrophages may modify CNS circuits that regulate locomotion remains an open question. Because brain infiltration and increased circulation of neutrophils were attributed to the systemic inflammation in the developing *nlrc3l* mutant embryo^28^, we sought to determine whether neutrophils are abnormally trafficked to the mutant larval brain. Consistent with this, neutrophils were found to be both residing in and moving through the diencephalon and other brain regions at 5-6 dpf, notably through the brain ventral vasculature surrounding the diencephalic choroid plexus (cp) and in close contact with diencephalic neurons (Fig. 7a-d and Supplementary Movies 8-9). By contrast, as is known under normal conditions^96^, slow moving and infiltrating neutrophils in the brain are not found in heterozygous or wild-type siblings, only rare occasions of individual neutrophils that rapidly circulate through the brain^28^ (Fig. 7a-d, Supplementary Movie 10). Compared with control siblings, *nlrc3l* mutants had a significant and unusual presence of brain lingering neutrophils and an increased number of peripheral neutrophils (Fig. 7c) at the time of our locomotor tracking at 5-6 dpf. Furthermore, the interaction of mutant neutrophils with the CNS begins early during development by 2 dpf (Fig. 7d and Supplementary Movies 11).

**Figure 7.**
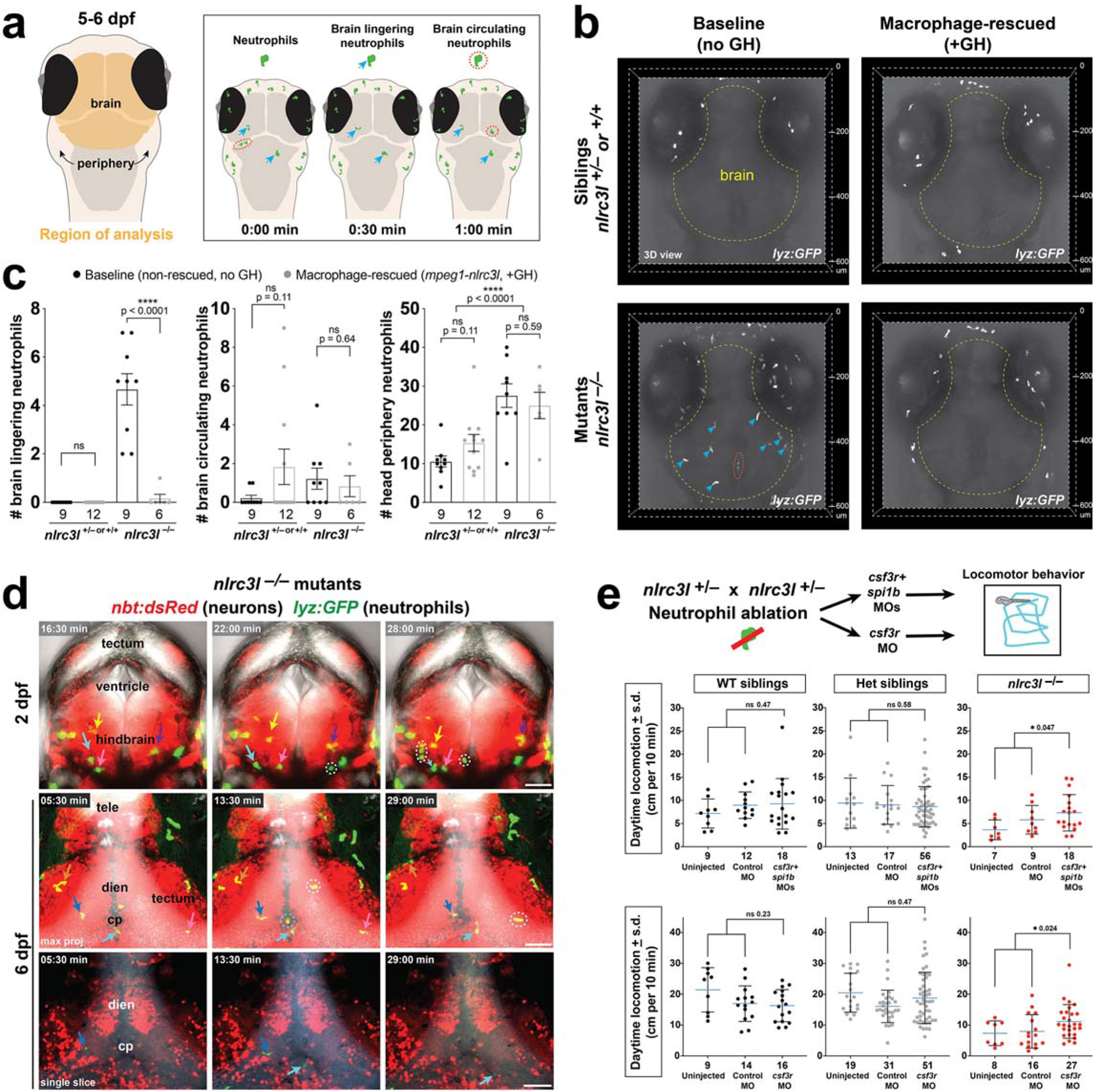
Reduced locomotion in *nlrc3l* mutants coincides with macrophage-dependent neutrophil infiltration of brain, and is partially reversed by neutrophil ablation. (a) Schematic showing brain region and categories of neutrophils analyzed in 5-6 dpf larvae from *nlrc3l* heterozygous intercross with (+GH) and without (no GH) the macrophage rescue transgene (*mpeg1-nlrc3l*). Lingering (blue arrow) defines infiltrated neutrophils that remain in the brain over the entire 15-minute period of imaging, while circulating (red dotted circle) defines neutrophils observed only at a single timepoint (imaged every 30-second) that flow through the brain. GH, GFP+ heart. (b) Representative 3D volumetric images of neutrophils (*lyz:GFP*+) in the brain at baseline and after macrophage rescue. See associated Supplementary Movies 9, 10, 12, and 13 showing the same zebrafish larvae as shown here. (c) Scatter bar plots show quantifications of brain lingering and circulating neutrophils, and total neutrophil numbers in the head periphery. Macrophage rescue reversed neutrophil infiltration in the *nlrc3l* larval brain, but not an overall increase in neutrophil numbers. Error bars show standard error of mean. Significance was determined by two-tailed t-tests. (d) Still images extracted from time-lapse confocal imaging showing circulating and infiltrating neutrophils in the brain at embryonic stage of 2 dpf and larval stage of 6 dpf (see associated Supplementary Movies 6 and 7). Individual neutrophils lingering in the parenchyma are labeled by arrows in different colors over three timepoints. Dotted circles mark circulating neutrophils passing through with blood flow. tele, telenchephalon; dien, diencephalon; cp, chorid plexus; max proj, maximum projection of a z-stack. (e) Genetic ablation of neutrophils using combined morpholinos against *csf3r* and *spi1b* and single morpholino against *csf3r* (top diagram). Uninjected and control morpholino injected groups show the expected reduced daytime locomotion in *nlrc3l* mutants. By contrast, daytime locomotion was significantly improved in *nlrc3l* mutants after neutrophil ablation using both morpholino strategies. No effect on locomotion was found in wild-type and heterozygous siblings after morpholino-mediated neutrophil ablation compared with baseline controls. Scatter plots show mean ± standard deviation from data of individual animals. Two-tailed t-tests were used to determine statistical significance. *n*, number of animals analyzed shown below each bar graph. See also associated Supplementary Movies 8-13.

The significant presence of neutrophils in the larval brain raises the possibility that inflammatory macrophages may indirectly modulate CNS circuits through their interaction with neutrophils in *nlrc3l* mutants. To examine the impact of macrophages on neutrophil activity in the larval *nlrc3l* mutants, we determined whether infiltration of neutrophils into the brain was dependent on *nlrc3l* function in macrophages, or conversely in neutrophils or both cell types. We generated stable transgenic lines to express wild-type *nlrc3l* in a macrophage-specific (Fig. 7a-d) or neutrophil-specific (Supplementary Fig. 10) manner in *nlrc3l* mutants and analyzed lingering, circulating, and total peripheral neutrophil numbers in or around the brain using in vivo time-lapse confocal imaging (Supplementary Movies 9-10 and 12-13). We found that a neutrophil-specific rescue of *nlrc3l* function in *nlrc3l* mutants did not reverse either the brain lingering neutrophil or reduced locomotor phenotype (Supplementary Fig. 10), whereas restoring wild-type *nlrc3l* expression in macrophages led to rescue of these phenotypes (Figs. 3 and 7, Supplementary Fig. 3, Supplementary Movies 9-10 and 12-13). The data together indicate that *nlrc3l* has non-cell-autonomous functions in affecting neutrophils and their impact on locomotor behavior through its action in macrophages.

We next tested the possibility that activated neutrophils in *nlrc3l* mutants may directly interfere with the locomotor circuit by using well established morpholino combination-based gene knockdowns against the granulocyte colony-stimulating factor receptor G-CSFR (gene name of *csf3r*) and myeloid transcription factor PU.1 (gene name of *spi1b*) that effectively removes myeloid cells^97, 98^, or against *csf3r* alone, which reduces neutrophil numbers but not macrophages as *pu.1/spi1b* knockdown does^71^, and assessed subsequent locomotor behaviors (Fig. 7e). We confirmed the high efficacy of the morpholino approaches to deplete neutrophils (90% average reduction)(Supplementary Fig. 11). Using the large-scale 96-well locomotor assay over the normal 14-hour day and 10-hour night cycles, we compared routine daytime swimming in *nlrc3l* mutants with their wild-type and heterozygous siblings after neutrophil ablation by treatment of *csf3r/spi1b* morpholinos (MOs) or single *csf3r* MO, and in control conditions (uninjected and control standard morpholino injected) on the same plate for each experiment (Fig. 7e). We found that in contrast to baseline or control morpholino-injected *nlrc3l* mutants, *nlrc3l* mutants after neutrophil depletion recovered significant swimming levels that are closer or similar to that of control siblings (Fig. 7e), indicating an essential role for neutrophils in causing the locomotor deficit. Furthermore, the effect of the morpholino treatments was specific to *nlrc3l* mutants, as neither wild-type nor heterozygous individuals had significantly altered locomotor behaviors (Fig. 7e). These results further substantiate the possibility that instead of direct signaling to neurons in the brain locomotor center, peripheral inflammatory macrophages activate neutrophils to infiltrate the brain to enable neutrophils to locally interact with and alter the locomotor circuits in *nlrc3l* mutants.

## Discussion

### Immune modulation of brain circuitry controlling innate spontaneous locomotion

While the wiring of the nervous system for implementing sensation and perception such as in vision, audition, olfaction, gustation, and touch has received much attention^99, 100^, the neural basis of unprovoked spontaneous behaviors remains underexplored. Spontaneous or exploratory locomotion of animals at homeostasis may reflect some degree of motivation in response to internal or external cues for survival; these animals may be moving to explore, forage for food, or find a mate^35–37, 59, 60^. Since changes to the immune system have been implicated in sickness and abnormal social behaviors characterized by reduced physical activity and motivation^1, 5^, we sought to determine whether dysregulated macrophages may alter brain circuits controlling normal locomotor activity. We leveraged investigation of the *nlrc3l* loss-of-function mutants as it provided an effective genetic paradigm for assessing the impact of inappropriate macrophage inflammation on brain circuit control of locomotion.

Using a large-scale and blinded platform to perform long-term tracking of spontaneous movements, we show that *nlrc3l* mutant larval zebrafish, which have aberrant inflammatory macrophages and no microglia, exhibit reduced frequency of exploratory swimming, yet maintain normal patterns of swimming (Fig. 2). When they are subjected to a series of light-to-dark visual stimuli, the mutants are fully capable of swimming at the same frequency and pattern as their wild type and heterozygous siblings, indicating no overt physical disability but seemingly decreased motivation for locomotor activity (Supplementary Fig. 2). In fact, after a series of normal locomotor responses to the visual stimuli, these mutants when placed back into their normal day-night cycles revert to displaying reduced daytime swimming (Supplementary Fig. 2). These results implicate that inappropriately activated peripheral macrophages during development or over a prolonged period are capable of perturbing brain locomotor circuits at the high level where the decision to move is made, rather than the downstream execution of the body movements as conveyed in our working model (Fig. 8). The precise aspects of the neural circuits controlling spontaneous swimming that may be modulated by pro-inflammatory innate immune functions remain to be determined by dissection and manipulation of the relevant circuitry.

**Figure 8.**
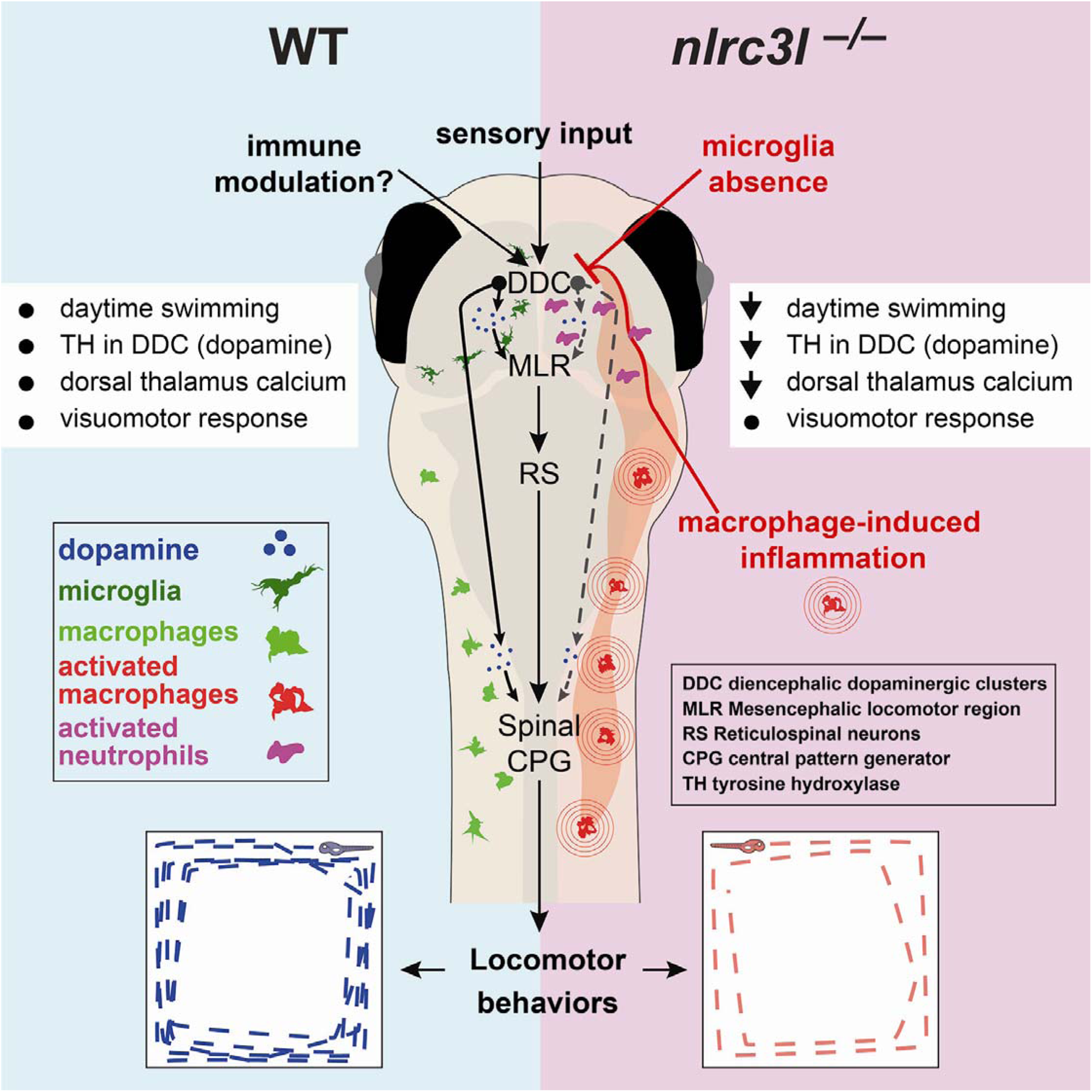
A working model for immunomodulation of daytime locomotor circuit in the larval zebrafish. Comparison between wild-type and *nlrc3l* mutant zebrafish conveys our current model. Sensory transduction based on a visuomotor response remains intact in *nlrc3l* mutants, while we show that immunomodulation is an important factor regulating larval zebrafish locomotion. The overall results suggest an inhibitory effect from macrophage-induced inflammation compounded by microglia loss on dopamine neurotransmission and subsequent activation of the descending neural circuits executing daytime locomotion. The neural circuits underlying locomotor drive may constitute diencephalic dopaminergic clusters (DDC) that either directly activate the spinal circuits (that include the central pattern generator (CPG)), or indirectly activate the mesencephalic locomotor region (MLR) and reticulospinal neurons (RS) to execute or potentiate swimming. Macrophage-induced inflammation activates neutrophils to infiltrate several regions of the brain, most notably in the vicinity of the diencephalic dopaminergic neurons where they may locally modulate neural circuits that control locomotor drive. List of effects: black dot indicates normal/typical status, and downward arrow means a reduction. Reversal of the locomotor deficit in *nlrc3l* mutants by restoring wild-type macrophages indicates that the behavioral phenotype is driven by mutant inflammatory macrophages. Further optogenetic or chemogenetic manipulations of the proposed neural circuit are needed to validate the working model.

### Inflammatory macrophages as a major cause of reduced locomotor drive in *nlrc3l* mutants

We show that the reduced locomotor drive phenotype in *nlrc3l* mutants has a myeloid origin. Using stable transgenics to restore wild-type expression of *nlrc3l* in a macrophage-specific manner in *nlrc3l* mutants, the locomotor deficit was significantly reversed, pointing to mutant macrophages as a major source of the behavioral phenotype. Since these mutants lack microglia and only have macrophages in the periphery, the mechanism by which the mutant macrophages could interfere with brain circuit function is expected to be indirect or mediated by soluble factors. Consistent with this, reversing infiltration of neutrophils in the brain by neutrophil ablation, restoring macrophage *nlrc3l* function, or inhibiting inflammatory signaling by dexamethasone ^101, 102^, as distinct means of suppressing inflammation were sufficient to significantly restore normal locomotor activity in *nlrc3l* mutants (Fig. 7 and Supplementary Figs. 4 and 10). Taken together, we show that inflammatory macrophages may indirectly, through yet unknown cues that may involve cytokines and neuromodulating molecules or recruitment of other leukocytes, alter the brain neural circuits regulating spontaneous behaviors at homeostasis (Fig. 8).

### Trafficking of neutrophils as an effective means for peripheral immune system to modulate CNS circuit functions

The view that the central nervous system (CNS) is immune privileged and immunologically inert and separated from the peripheral immune system has been revised in recent decades^18, 21^. Under normal conditions, neutrophils are rarely found in the CNS as they are restricted from trafficking into the brain parenchyma and ventricular regions due to the blood-brain barrier^96, 103^. However, upon a brain perturbation or pathology, such as trauma, infection, ischemia, neurodegeneration or autoimmunity, neutrophils are recruited into the brain and found to be lingering in the CNS environment^96, 103^. Interestingly, while we found the expected absence of infiltrating neutrophils in wild-type and heterozygous larval zebrafish brain, the presence of infiltrating and lingering neutrophils in the brain of *nlrc3l* mutants was striking (Fig. 7 and Supplementary Movies 9-10). Beginning at embryonic development and continuing through mature larval stages, *nlrc3l* mutants exhibit the unusual activities of neutrophils passing through and lingering in the brain, as shown previously^28^ and in this study (Fig. 7 and Supplementary Movie 9), a phenomenon associated with inflammatory brain conditions^96, 103^. The neutrophil infiltration into the larval brain is secondary to the inappropriate activation of macrophages in *nlrc3l* mutants as it can be reversed by restoring wild-type macrophages but not by wild-type neutrophils in *nlrc3l* mutants (Fig. 7 and Supplementary Fig. 10), indicating a requirement for the mutant macrophages. These results indicate that the effects of the *nlrc3l* mutation on neutrophils were non-cell-autonomous (Supplementary Fig. 10). While the precise nature of the neutrophil interaction with the CNS neurons in *nlrc3l* mutants is yet unknown, it is possible that known neutrophil production of effector molecules (such as free oxygen radicals and extracellular traps composed of proteins and DNA called NETs)^96, 103^ may impact the surrounding neurons to affect locomotor control in the diencephalon. We further show that depletion of neutrophils by a combined morpholino-mediated knockdown of *csf3r* and *spi1b,* or single *csf3r* knockdown was sufficient to partially restore normal locomotion in *nlrc3l* mutants, providing evidence that modulation of the locomotor circuitry in the ventral larval zebrafish brain was in part mediated by activated neutrophils in the *nlrc3l* mutants (Fig. 7e). Partial recovery of the behavioral deficit may be due to incomplete neutrophil ablation, or other neutrophil-independent factors that disrupt normal behavioral circuit function stemming from other mutant cell types, in particular but not exclusively macrophages. Collectively, these results indicate that the inflammatory macrophages trigger neutrophil infiltration into the brain, allowing activated neutrophils to directly interact with the brain circuitry that controls innate spontaneous locomotion (Fig. 8).

### Microglia loss has negligible impact on spontaneous locomotion

Besides macrophages in the periphery, we also examined the role of microglia, which are brain-resident macrophages that have been widely anticipated and implicated to affect CNS circuitry and functions due to their role in synaptic pruning and surveillance during development^17, 65, 66, 104^. Microglia are absent in *nlrc3l* mutants, because they are derived from primitive macrophages, which are inappropriately activated and thereby diverted from their normal developmental course ^28^. In order to delineate the causal factor(s) leading to the behavioral deficit in *nlrc3l* mutants, we investigated three other microglia-lacking mutants, gene knockouts of *irf8*, *xpr1b, and pu.1/spi1b*, to determine whether a sole microglia loss without the inflammatory phenotype could result in reduced daytime locomotion. Our results revealed minimal to no effect of microglia loss on locomotor behavioral change in *irf8*, *xpr1b, and pu.1/spi1b* mutants (Fig. 4c-g).

Our finding that an absence of microglia in larval zebrafish at 5-8 dpf did not cause a significant disruption to routine swimming behaviors seems consistent with murine studies indicating no significant behavioral or cognitive impairment with removal of microglia at baseline^105^. Understanding the full extent of microglia impact on behavioral outcomes remains incomplete, while murine studies raise the possibility that microglia may be primary responders to homeostatic perturbations in the CNS, rather than main drivers of CNS maintenance and homeostasis^21, 105–108^. Furthermore, in contexts of CNS pathology or inflammation, a role for microglia in modifying CNS circuits and behavior remains supported and possible^21, 104^. Consistent with this, our study underscores the possibility that systemic inflammation stemming from dysregulated peripheral macrophages intensifies the effects of microglia loss, thereby causing reduced locomotion and sickness-like behavior. Microglia may normally have a role in dampening inflammatory effects in the CNS, thereby limiting potentially negative effects on circuit functions and behavioral outcomes. Similarly, *nlrc3l* mutants may be more severely impacted by the chronic systemic inflammation due to an absence of microglia which normally limit pro-inflammatory signaling in the CNS.

### Dopaminergic control of spontaneous locomotion may be susceptible to immune modulation

We found that the reduced locomotion in *nlrc3l* mutants was associated with a disruption of the diencephalic dopaminergic circuit previously known to control swimming^34, 69, 86, 87^ (Fig. 8). *nlrc3l* mutants at larval stages were found to have reduced intracellular calcium in the dorsal thalamus and DDC region at baseline (Fig. 5 and Supplementary Fig. 8) and a significant reduction in the rate-limiting enzyme TH for dopamine synthesis in the diencephalic dopaminergic clusters (DDC) (Fig. 6). These results are consistent with the known importance of dopamine neurotransmission and brain dopaminergic circuits for initiation and control of locomotion that is evolutionarily conserved from fish to humans^109–112^. DDC encompass neurons that directly control spinal motor neurons essential for swimming^69^ and provide the primary source of dopamine to the spinal cord neurons including the central pattern generator (CPG) that control body movements^30, 34, 86^. We therefore propose that appropriate levels of dopamine neurotransmission from the diencephalon is essential to regulate locomotion either directly from the DDC to the spinal CPG or via the mesencephalic locomotor region (MLR) to the reticulospinal neurons (RS) in the hindbrain (Fig. 8) known to convey motor commands from the brain to the spinal circuits in zebrafish^69, 90, 113–115^. This study implicates that the dopaminergic circuits underlying locomotor control may be more susceptible or responsive to inflammatory signaling, thereby subject to innate immune modulation, although the precise mechanism remains to be determined.

In summary, this study leverages genetic and large-scale approaches in zebrafish using *nlrc3l* and a set of microglia-lacking mutants to reveal the capability of peripheral inflammatory macrophages in the absence of microglia to alter function of the locomotion-controlling brain circuits through recruitment of neutrophils to the brain. By contrast, we found that microglia loss alone had negligible consequence on spontaneous locomotor behaviors. Overall, we reveal novel interactions by which chronic peripheral inflammation evoked by inflammatory macrophages can intercept central neural circuits through co-opting neutrophils to infiltrate the brain and lead to spontaneous locomotor behavioral changes.

## Materials and Methods

### Zebrafish

Embryos from wild-type (TL and AB), mutant and transgenic backgrounds: *nlrc3l^st73^* ^28^, *irf8^st95^* ^48^, *xpr1b^st89^* ^49^, *pu.1/spi1b^fh509^* ^50^, *mpeg1:EGFP* ^116^, *lyz:GFP* ^117^, *neurod:GCaMP6f* ^80^, and *nbt:DsRed* ^118^ were raised at 28.5°C and staged as described ^119^. Stable transgenic *mpeg1:nlrc3l* line was generated using Tol2-mediated transgenesis based on a previously cloned construct ^28^ combining full-length wild-type *nlrc3l* coding sequence downstream of *mpeg1(1.86 kb)*^116^ regulatory sequence. Stable *elavl3:GCaMP6s* line was generated using Tol2-mediated transgenesis. This study was carried out in accordance with the approval of UNC-Chapel Hill Institutional Animal Care and Use Committee (protocols 16–160 and 19–132).

### Whole mount RNA in situ hybridization

RNA in situ was performed using standard methods. Antisense riboprobes for *mfap4*, *mpx*, and *irg1* were synthesized as described ^28, 71, 120^. Additional antisense riboprobes were synthesized using primers targeting coding sequences of *dbh*, *gad1b*, *nr4a2b/nurr1*, *serta/slc6a4a*, and *vglut2a/slc17a6b* (sequences detailed in Supplementary File 1).

### RNA isolation and qPCR

RNA was isolated using the RNAqueous-Micro kit RNA Isolation Procedure (Ambion). Whole larvae were lysed in 100–300 uL RNA lysis buffer. cDNA was made from 150 or 200 ng of total RNA using oligo (dT) primer with SuperScript IV reverse transcriptase (Invitrogen). qPCR was performed on the QuantStudio 3 Real-Time PCR System (Applied Biosystems) using SYBR Green. The delta-delta ct method was used to determine the relative levels of mRNA expression between experimental samples and control. *ef1a* was used as the reference gene for determination of relative expression of all target genes. Primer sequences for qPCR analysis are listed in Supplementary file 1. All qPCR experiments were conducted using 3 technical replicates and a minimum of 3 biological replicates.

### RNA-seq analysis

Paired-end RNA-Sequencing was performed on two independent biological replicates of 10 or 12 pooled 4-dpf larval zebrafish per genotypic group (*nlrc3l* mutants versus their siblings) by Illumina HiSeq 2×100 basepairs. The mutants were pre-sorted by neutral red for lack of microglia while the siblings, consisting of heterozygotes and wild-types, were sorted for presence of normal microglia. Genotypes were verified using PCR-restriction enzyme assay prior to library preparation. RNA-seq analysis followed previously developed protocol^121^, which consists of modules and packages including FastQC for quality control, Trimmomatic for data trimming, and HISAT2 for sequence alignment. The differential expression analysis was performed using DESeq2. Gene ontology analysis was performed on the differentially expressed genes with bioMaRt in R. All RNA-seq data has been submitted to the GEO repository to be accessed under the accession GSE179883.

### In vivo time-lapse and static confocal imaging

All time-lapse and static z-stack imaging were performed using a Nikon A1R+ hybrid galvano and resonant scanning confocal system equipped with an ultra-high speed A1-SHR scan head and controller. Images were obtained using an apochromat lambda 40x water immersion objective (NA 1.15) or a plan apochromat lambda 20x objective (NA 0.75). Z-steps at 1–2 µm were taken at 40x and 3–5 µm at 20x. Different stages of zebrafish were mounted on glass-bottom dishes using 1.5% low-melting agarose and submerged in fish water supplemented with 0.003% PTU to inhibit pigmentation. All image acquisition parameters (including laser power, gain, scan mode/speed, and resolution) were set constant for experiments requiring fluorescence quantification. Raw data was used for quantification, and brightness/contrast adjustments may be independently applied to whole images only for the purpose of clarity in figure presentation.

### 96-well or 24-well behavior experiment setup

Heterozygous intercrosses were conducted to generate homozygous mutants that were co-housed and monitored in identical conditions as their heterozygous and wild-type siblings. Embryos were raised with daily water change and health checks at a density of 50-100 per dish at 28.5 _C_ without the light:dark cycles until they reached 5 dpf. Fish were checked for normal gross morphology including an inflated large swim bladder and normal motility before and after behavioral tracking. Although infrequent, any larvae that appeared sick or have deflated swim bladder were eliminated from further analyses. Individual 5 dpf larvae were placed alone in each well of a 96-well (Fig. 2a) or 24-well plate (Fig. 3a, Supplementary Fig. 2), and simultaneously tracked in the customized ZebraBox built by ViewPoint Behavior Technology. At 5 dpf we are able to avoid possible confounding effects from feeding since the larval zebrafish can survive on their yolk contents. The automated tracking system recorded the location of every animal on the plate continuously using an ultrasensitive infrared CCD camera. Because larvae undergo spontaneous burst and glide swimming interspersed with short periods of inactivity, data was automatically collected at 10-minute integrations^43^ for 72-hour tracking under 14-hour light: 10-hour dark cycles (432 total timepoints)(Figs. 2, 3), or 1-minute integrations for the 6-hour light-dark visuomotor assay (360 total timepoints)(Supplementary. Fig. 1). The larvae were housed in the ZebraBox at room temperature to maintain them at around 6 dpf so the impact of development is minimal while preserving their active and healthy state. 96 deep well polypropylene plates (0.7 ml) with flat bottom and square wells (EK-2074, E&K Scientific; 201242-100, Agilent), or 24 well cell culture clear plates with flat bottom and circular wells (T1024, Thomas Scientific) were used. For 72-hour experiments, water evaporation was minimal in the 24-well setup but significant in the 96-well plate, which required a daily replenishment of 50-100 ul of fish water per well. Animals were genotyped after completion of behavioral tracking to ensure unbiased data collection.

### Large-scale locomotor behavior analysis

High-throughput locomotor behavior data collected was mined and quantified in Matlab and R. The outputs from ZebraBox for each experiment include a spreadsheet of raw data for all animals at all timepoints by variables (such as counts, distance, duration of movements) and the images of movement traces, which were integrated over a 10-minute window for the entire 3-day duration of tracking. Extraction of features such as distance and duration of movements at time points and periods of interest was achieved in Matlab. Key locomotor metrics (distance, frequency, and speed) for each individual fish were determined as an average across time periods of interest (such as d2 and d3 light periods for 72-hour tracking, unless otherwise noted). We did not include the first day given its variability during acclimation.

### Locomotor pattern analysis

Matlab was used to process images of movement traces recorded at every timepoint corresponding to daytime d2 and d3 periods to digitally segment each well into 9 equal squares in a 3 x 3 grid to determine pixel values for each square of the grid for all fish at all timepoints of interest (Fig. 3g). R program was used to compute the perimeter and center scores from the outputs of the pixel-based pattern analysis in Matlab. The perimeter swimming was quantified using all but the center square for each individual at every timepoint; each outer square was assigned 0 for pixel value _<_ 9 (baseline no movement) or 1 for pixel value > 9, then these numbers were summed to generate a single perimeter score per individual per time point, ranging from 0 (no perimeter swimming) to 8 (full perimeter swimming) (Fig. 3g). The center square was evaluated directly by its pixel value (<100 for little to no center activity and >550 for high center activity). Downstream statistical analyses of the quantified locomotor patterns, such as hierarchical clustering, were performed and visualized using various R packages, including heatmap.2 and hclust, or GraphPad Prism 9.

### Visuomotor assay using light-dark conditions

For the visuomotor assay, the animals were subjected to a series of defined timed intervals of alternating 100% light or 0% light (for dark) conditions implemented by the built-in light control program in the Zebrabox and locomotor behaviors were recorded every minute for a total tracking time of 6 hours (Supplementary Fig. 1). We programmed our system light control to begin with a 3-hour full lighting for an acclimation period followed by alternating dark and light periods dispersed over four 30-minute intervals and six 10-minute intervals to allow assessment of 5 repeated light-to-dark switches (Supplementary Fig. 1). For analysis, we extracted data from specific periods of response to visual stimuli, and calculated locomotor metrics for each individual animal.

### *In vivo* brain calcium imaging to measure spontaneous daytime neural activity

Double transgenic larval zebrafish carrying *neurod:GCaMP6f* and *nbt:dsRed* at 5 dpf were acclimated for ∼ 24 hours in 14h light: 10h dark cycle prior to brain imaging at 6 dpf. Two z-planes (tectum at ∼50 microns below skin and diencephalon at ∼ 55 microns below tectum slice) were imaged at 1 scan per second for 10 minutes in ambient light using resonant scanning at 16X line integration on a Nikon A1R+ confocal microscope with a 40x water immersion objective (NA 1.15). Genotyping ensued imaging. *GCaMP6f* fluorescence was extracted for ROIs in the telencephalon, diencephalon, tectum and cerebellum over the entire imaging period using ImageJ. R program was used to calculate spontaneous neural activity of the ROIs by ΔF/F_0_, either from discrete regions of interest or random sampling of individual neurons. Baseline calcium (F_0_) of each ROI was determined by the average fluorescence intensity over all time points. Number of calcium spikes was determined using Matlab by setting a threshold ΔF/F_0_ value for each brain region ROIs to define the number of discrete events at which ΔF/F_0_ is above the set threshold with at least one timepoint gap from the next spike event. Relative somatic calcium fluorescence for each ROI was calculated as an average of the relative changes in *GCaMP6f* fluorescence from baseline fluorescence over all timepoints of the 10-minute calcium imaging per fish. Due to the varying *neurod:GCaMP6f* reporter expression intrinsic to Tol2-mediated transgenesis and random sampling, we found *GCaMP6f* signals consistently strong in the telencephalon but not in the diencephalon that already has low baseline fluorescence. Animals with *GCaMP6f* signals that cannot be delineated in at least a subpopulation of the diencephalic neurons were excluded from diencephalon analysis. Pan-neuronal *elavl3:GCaMP6s* transgenic zebrafish were acclimated, imaged at 1 Hz for a minimum of 5 minutes in ambient light using confocal imaging with a 40x objective, and analyzed using the same pipeline as above.

### Small-molecule anti-inflammatory drug treatment

Administration of different small-molecule drugs was conducted by adding each individual chemical into the fish water starting at ∼24 hours prior to behavioral tracking at 5 dpf. Fish water freshly supplemented with the chemical was continuously used in the behavioral plate in which the larvae were individually housed in each well. Daily replenishment of chemical-supplemented water of about 50-100 ul was applied to refill the wells in the 96-well format plate due to evaporation. Bay 11–7082 (1 µM), dexamethasone (3.25 µM), and 17-DMAG (5 µM) were used as listed in Supplementary file 1. Behavioral tracking ensued for 72 hours in normal day-night cycles.

### Gene knockdown by morpholino microinjection

Morpholino oligomers were injected into one-cell stage embryos from *nlrc3l* heterozygous intercrosses. Antisense morpholino oligos were purchased from Gene Tools and re-suspended in water to make 1-3 mM stocks. Morpholino sequences are listed in Supplementary file 1. Morpholinos were heated at 65°C for 5 min and cooled to room temperature before microinjection at 1 nL. Efficacy of the morpholinos were validated based on the known phenotype. Quantification of neutrophils at 2 dpf was based on a cell count and at 5-6 dpf based on total area of neutrophils in a ROI using ImageJ. Area was used at later stages due to an increased presence of clustering and interconnected neutrophils, which makes cell counting less reliable.

### Neutral red staining

Microglia were examined in live larvae by neutral red vital dye staining as previously described^28, 48^. Larvae at 3-4 dpf were stained by immersion in fish water supplemented with 2.5 μg/mL neutral red and 0.003% PTU at 28.5°C for 1 hr, followed by 1–2 water changes, and then analyzed using a stereomicroscope.

### TH immunostaining

Standard whole mount tissue immunostaining was followed using the rabbit anti-Tyrosine Hydroxylase antibody (AB152, Sigma Aldrich) at 1:100 dilution. Briefly, zebrafish larvae were fixed overnight in 4% paraformaldehyde followed by methanol permeabilization, and incubated with primary antibody in blocking solution overnight at 4°C and then secondary antibody, goat anti-rabbit Alexa Fluor 594 (ThermoFisher), at 1:3000 dilution in blocking solution for detection.

### Statistical analyses

Unpaired two-tailed t-tests were performed unless otherwise noted. F test was used to compare variances. For unequal variances, Welch’s correction was used on the two-tailed t-test. For multiple comparisons of 3 or more groups, one-way ANOVA test was applied followed by multiple pair-wise comparisons to determine the pair(s) showing significant differences using FDR-adjusted p-values. All graphical plots and statistical tests were generated using GraphPad Prism 9 unless otherwise noted. All box-whisker plots show the median, first and third quartiles, minimum, and maximum of the data plotted.

## Author contributions

CES: conceptualization, data analysis and visualization, supervision, funding acquisition, behavior experiments and computational analyses, generation of rescue transgenic line, writing-original draft; VK: experimentation (behavior, RNA in situ, imaging, immunohistochemistry, genotyping, morpholinos, qPCR) and data analysis; PC: computational pipeline for behavior analyses and RNA-seq with CES, Matlab and R scripts, data analysis; CTD: experimentation (behavior, RNA in situ, calcium and confocal imaging, genotyping, qPCR); VH: experimentation (behavior, imaging, genotyping, qPCR); CGS: experimentation and analysis of calcium imaging; AMR: experimentation (macrophage rescue characterization, qPCR) and all schematic illustrations. All authors reviewed and edited the manuscript.

## Acknowledgements

We thank Ana Meireles, William Talbot, John Rawls, Cecilia Moens and Claire Wyart for the generous sharing of fish lines and reagents. We are grateful for the substantial insights and discussions from Jonathan Gable on our manuscript. Michelle Altemara and her staff at the UNC Zebrafish Aquaculture Core Facility conducted critical support for zebrafish housing and care. Pilot behavior experiments prior to this study benefited from critical insights and inputs given by Constance Richter and Alex Schier. This work was supported by NIH NIGMS grant 1R35GM124719 to C.E.S.

## References

1. Kirsten, K., Soares, S.M., Koakoski, G., Carlos Kreutz, L. & Barcellos, L.J.G. Characterization of sickness behavior in zebrafish. Brain Behav Immun 73, 596–602 (2018).

2. Dantzer, R., O’Connor, J.C., Freund, G.G., Johnson, R.W. & Kelley, K.W. From inflammation to sickness and depression: when the immune system subjugates the brain. Nat Rev Neurosci 9, 46–56 (2008).

3. Kelley, K.W. et al. Cytokine-induced sickness behavior. Brain Behav Immun 17 Suppl 1, S112–118 (2003).

4. Hart, B.L. Biological basis of the behavior of sick animals. Neurosci Biobehav Rev 12, 123–137 (1988).

5. Dantzer, R., Heijnen, C.J., Kavelaars, A., Laye, S. & Capuron, L. The neuroimmune basis of fatigue. Trends Neurosci 37, 39–46 (2014).

6. Kulkarni, O.P., Lichtnekert, J., Anders, H.J. & Mulay, S.R. The Immune System in Tissue Environments Regaining Homeostasis after Injury: Is “Inflammation” Always Inflammation? Mediators Inflamm 2016, 2856213 (2016).

7. Nathan, C. Points of control in inflammation. Nature 420, 846–852 (2002).

8. Dunn, A.J. Effects of cytokines and infections on brain neurochemistry. Clin Neurosci Res 6, 52–68 (2006).

9. Dantzer, R. Cytokine, sickness behavior, and depression. Immunol Allergy Clin North Am 29, 247–264 (2009).

10. Bluthe, R.M., Michaud, B., Poli, V. & Dantzer, R. Role of IL-6 in cytokine-induced sickness behavior: a study with IL-6 deficient mice. Physiol Behav 70, 367–373 (2000).

11. Bluthe, R.M. et al. Synergy between tumor necrosis factor alpha and interleukin-1 in the induction of sickness behavior in mice. Psychoneuroendocrinology 19, 197–207 (1994).

12. Oishi, Y. & Manabe, I. Macrophages in inflammation, repair and regeneration. Int Immunol 30, 511–528 (2018).

13. Watanabe, S., Alexander, M., Misharin, A.V. & Budinger, G.R.S. The role of macrophages in the resolution of inflammation. J Clin Invest 129, 2619–2628 (2019).

14. Torraca, V., Masud, S., Spaink, H.P. & Meijer, A.H. Macrophage-pathogen interactions in infectious diseases: new therapeutic insights from the zebrafish host model. Dis Model Mech 7, 785–797 (2014).

15. Schultze, J.L., Schmieder, A. & Goerdt, S. Macrophage activation in human diseases. Semin Immunol 27, 249–256 (2015).

16. Chiot, A. et al. Modifying macrophages at the periphery has the capacity to change microglial reactivity and to extend ALS survival. Nat Neurosci 23, 1339–1351 (2020).

17. Wright-Jin, E.C. & Gutmann, D.H. Microglia as Dynamic Cellular Mediators of Brain Function. Trends Mol Med 25, 967–979 (2019).

18. Papadopoulos, Z., Herz, J. & Kipnis, J. Meningeal Lymphatics: From Anatomy to Central Nervous System Immune Surveillance. J Immunol 204, 286–293 (2020).

19. Kipnis, J. Multifaceted interactions between adaptive immunity and the central nervous system. Science 353, 766–771 (2016).

20. Filiano, A.J., Gadani, S.P. & Kipnis, J. How and why do T cells and their derived cytokines affect the injured and healthy brain? Nat Rev Neurosci 18, 375–384 (2017).

21. Norris, G.T. & Kipnis, J. Immune cells and CNS physiology: Microglia and beyond. J Exp Med 216, 60–70 (2019).

22. Alves de Lima, K., Rustenhoven, J. & Kipnis, J. Meningeal Immunity and Its Function in Maintenance of the Central Nervous System in Health and Disease. Annu Rev Immunol 38, 597–620 (2020).

23. Frenois, F. et al. Lipopolysaccharide induces delayed FosB/DeltaFosB immunostaining within the mouse extended amygdala, hippocampus and hypothalamus, that parallel the expression of depressive-like behavior. Psychoneuroendocrinology 32, 516–531 (2007).

24. Haba, R. et al. Lipopolysaccharide affects exploratory behaviors toward novel objects by impairing cognition and/or motivation in mice: Possible role of activation of the central amygdala. Behav Brain Res 228, 423–431 (2012).

25. Laye, S. et al. Endogenous brain IL-1 mediates LPS-induced anorexia and hypothalamic cytokine expression. Am J Physiol Regul Integr Comp Physiol 279, R93–98 (2000).

26. Qin, L. et al. Systemic LPS causes chronic neuroinflammation and progressive neurodegeneration. Glia 55, 453–462 (2007).

27. Campbell, I.L., Hofer, M.J. & Pagenstecher, A. Transgenic models for cytokine-induced neurological disease. Biochim Biophys Acta 1802, 903–917 (2010).

28. Shiau, C.E., Monk, K.R., Joo, W. & Talbot, W.S. An anti-inflammatory NOD-like receptor is required for microglia development. Cell Rep 5, 1342–1352 (2013).

29. Orger, M.B. & de Polavieja, G.G. Zebrafish Behavior: Opportunities and Challenges. Annu Rev Neurosci 40, 125–147 (2017).

30. Berg, E.M., Bjornfors, E.R., Pallucchi, I., Picton, L.D. & El Manira, A. Principles Governing Locomotion in Vertebrates: Lessons From Zebrafish. Front Neural Circuits 12, 73 (2018).

31. Grillner, S. & El Manira, A. Current Principles of Motor Control, with Special Reference to Vertebrate Locomotion. Physiol Rev 100, 271–320 (2020).

32. Schurger, A., Mylopoulos, M. & Rosenthal, D. Neural Antecedents of Spontaneous Voluntary Movement: A New Perspective. Trends Cogn Sci 20, 77–79 (2016).

33. Schurger, A., Sitt, J.D. & Dehaene, S. An accumulator model for spontaneous neural activity prior to self-initiated movement. Proc Natl Acad Sci U S A 109, E2904–2913 (2012).

34. Goulding, M. Circuits controlling vertebrate locomotion: moving in a new direction. Nat Rev Neurosci 10, 507–518 (2009).

35. Horstick, E.J., Mueller, T. & Burgess, H.A. Motivated state control in larval zebrafish: behavioral paradigms and anatomical substrates. J Neurogenet 30, 122–132 (2016).

36. Moscarello, J.M., Ben-Shahar, O. & Ettenberg, A. Effects of food deprivation on goal-directed behavior, spontaneous locomotion, and c-Fos immunoreactivity in the amygdala. Behav Brain Res 197, 9–15 (2009).

37. Stringer, C. et al. Spontaneous behaviors drive multidimensional, brainwide activity. Science 364, 255 (2019).

38. Teitelbaum P., S.T., Whishaw I.Q. Sources of Spontaneity in Motivated Behavior. Motivation. Springer, Boston, MA, 1983.

39. Yoder, J.A., Nielsen, M.E., Amemiya, C.T. & Litman, G.W. Zebrafish as an immunological model system. Microbes Infect 4, 1469–1478 (2002).

40. Santoriello, C. & Zon, L.I. Hooked! Modeling human disease in zebrafish. J Clin Invest 122, 2337–2343 (2012).

41. Howe, K. et al. The zebrafish reference genome sequence and its relationship to the human genome. Nature 496, 498–503 (2013).

42. Lam, S.H., Chua, H.L., Gong, Z., Lam, T.J. & Sin, Y.M. Development and maturation of the immune system in zebrafish, Danio rerio: a gene expression profiling, in situ hybridization and immunological study. Dev Comp Immunol 28, 9–28 (2004).

43. Rihel, J., Prober, D.A. & Schier, A.F. Monitoring sleep and arousal in zebrafish. Methods Cell Biol 100, 281–294 (2010).

44. Budick, S.A. & O’Malley, D.M. Locomotor repertoire of the larval zebrafish: swimming, turning and prey capture. J Exp Biol 203, 2565–2579 (2000).

45. Motta, V., Soares, F., Sun, T. & Philpott, D.J. NOD-like receptors: versatile cytosolic sentinels. Physiol Rev 95, 149–178 (2015).

46. Xu, J. et al. Temporal-Spatial Resolution Fate Mapping Reveals Distinct Origins for Embryonic and Adult Microglia in Zebrafish. Dev Cell 34, 632–641 (2015).

47. Ferrero, G. et al. Embryonic Microglia Derive from Primitive Macrophages and Are Replaced by cmyb-Dependent Definitive Microglia in Zebrafish. Cell Rep 24, 130–141 (2018).

48. Shiau, C.E., Kaufman, Z., Meireles, A.M. & Talbot, W.S. Differential requirement for irf8 in formation of embryonic and adult macrophages in zebrafish. PLoS One 10, e0117513 (2015).

49. Meireles, A.M. et al. The phosphate exporter xpr1b is required for differentiation of tissue-resident macrophages. Cell Rep 8, 1659–1667 (2014).

50. Roh-Johnson, M. et al. Macrophage-Dependent Cytoplasmic Transfer during Melanoma Invasion In Vivo. Dev Cell 43, 549–562 e546 (2017).

51. Wang, T. et al. Nlrc3-like is required for microglia maintenance in zebrafish. J Genet Genomics 46, 291–299 (2019).

52. Salem, S., Salem, D. & Gros, P. Role of IRF8 in immune cells functions, protection against infections, and susceptibility to inflammatory diseases. Hum Genet 139, 707–721 (2020).

53. Kuil, L.E. et al. Reverse genetic screen reveals that Il34 facilitates yolk sac macrophage distribution and seeding of the brain. Dis Model Mech 12 (2019).

54. Hall, C.J. et al. Immunoresponsive gene 1 augments bactericidal activity of macrophage-lineage cells by regulating beta-oxidation-dependent mitochondrial ROS production. Cell metabolism 18, 265–278 (2013).

55. Lampropoulou, V. et al. Itaconate Links Inhibition of Succinate Dehydrogenase with Macrophage Metabolic Remodeling and Regulation of Inflammation. Cell metabolism 24, 158–166 (2016).

56. Nemeth, B. et al. Abolition of mitochondrial substrate-level phosphorylation by itaconic acid produced by LPS-induced Irg1 expression in cells of murine macrophage lineage. FASEB journal : official publication of the Federation of American Societies for Experimental Biology 30, 286–300 (2016).

57. Sanderson, L.E. et al. An inducible transgene reports activation of macrophages in live zebrafish larvae. Developmental and comparative immunology 53, 63–69 (2015).

58. Hall, C.J., Sanderson, L.E., Crosier, K.E. & Crosier, P.S. Mitochondrial metabolism, reactive oxygen species, and macrophage function-fishing for insights. Journal of molecular medicine 92, 1119–1128 (2014).

59. Maye, A., Hsieh, C.H., Sugihara, G. & Brembs, B. Order in spontaneous behavior. PLoS One 2, e443 (2007).

60. Kobayashi, K., Shimizu, N., Matsushita, S. & Murata, T. The assessment of mouse spontaneous locomotor activity using motion picture. J Pharmacol Sci 143, 83–88 (2020).

61. Dunn, T.W. et al. Brain-wide mapping of neural activity controlling zebrafish exploratory locomotion. Elife 5, e12741 (2016).

62. MacPhail, R.C. et al. Locomotion in larval zebrafish: Influence of time of day, lighting and ethanol. Neurotoxicology 30, 52–58 (2009).

63. Burgess, H.A. & Granato, M. Modulation of locomotor activity in larval zebrafish during light adaptation. J Exp Biol 210, 2526–2539 (2007).

64. Peng, X. et al. Anxiety-related behavioral responses of pentylenetetrazole-treated zebrafish larvae to light-dark transitions. Pharmacol Biochem Behav 145, 55–65 (2016).

65. Prinz, M., Jung, S. & Priller, J. Microglia Biology: One Century of Evolving Concepts. Cell 179, 292–311 (2019).

66. Frost, J.L. & Schafer, D.P. Microglia: Architects of the Developing Nervous System. Trends Cell Biol 26, 587–597 (2016).

67. Kastenhuber, E., Kratochwil, C.F., Ryu, S., Schweitzer, J. & Driever, W. Genetic dissection of dopaminergic and noradrenergic contributions to catecholaminergic tracts in early larval zebrafish. J Comp Neurol 518, 439–458 (2010).

68. Lambert, A.M., Bonkowsky, J.L. & Masino, M.A. The conserved dopaminergic diencephalospinal tract mediates vertebrate locomotor development in zebrafish larvae. J Neurosci 32, 13488–13500 (2012).

69. Tay, T.L., Ronneberger, O., Ryu, S., Nitschke, R. & Driever, W. Comprehensive catecholaminergic projectome analysis reveals single-neuron integration of zebrafish ascending and descending dopaminergic systems. Nat Commun 2, 171 (2011).

70. Svahn, A.J. et al. Development of ramified microglia from early macrophages in the zebrafish optic tectum. Dev Neurobiol 73, 60–71 (2013).

71. Yang, L. et al. Drainage of inflammatory macromolecules from the brain to periphery targets the liver for macrophage infiltration. Elife 9 (2020).

72. Lee, J.-H. et al. Inhibition of NF-κB activation through targeting IκB kinase by celastrol, a quinone methide triterpenoid. Biochemical Pharmacology 72, 1311–1321 (2006).

73. Shimp, S.K. et al. HSP90 inhibition by 17-DMAG reduces inflammation in J774 macrophages through suppression of Akt and nuclear factor-κB pathways. Inflammation Research 61, 521–533 (2012).

74. Lee, J., Rhee, M.H., Kim, E. & Cho, J.Y. BAY 11-7082 is a broad-spectrum inhibitor with anti-inflammatory activity against multiple targets. Mediators of inflammation 2012, 416036 (2012).

75. Aghai, Z.H. et al. Dexamethasone suppresses expression of Nuclear Factor-kappaB in the cells of tracheobronchial lavage fluid in premature neonates with respiratory distress. Pediatric research 59, 811–815 (2006).

76. Rihel, J. et al. Zebrafish behavioral profiling links drugs to biological targets and rest/wake regulation. Science 327, 348–351 (2010).

77. Fotowat, H., Lee, C., Jun, J.J. & Maler, L. Neural activity in a hippocampus-like region of the teleost pallium is associated with active sensing and navigation. Elife 8 (2019).

78. Dumitrescu, A.S., Fidelin, K. & Wyart, C. Toward a comprehensive model of circuits underlying locomotion: What did we learn from zebrafish? Neural Control of Movement, 2020, pp 125–152.

79. Herbomel, P., Thisse, B. & Thisse, C. Zebrafish early macrophages colonize cephalic mesenchyme and developing brain, retina, and epidermis through a M-CSF receptor-dependent invasive process. Dev Biol 238, 274–288 (2001).

80. Rupprecht, P., Prendergast, A., Wyart, C. & Friedrich, R.W. Remote z-scanning with a macroscopic voice coil motor for fast 3D multiphoton laser scanning microscopy. Biomed Opt Express 7, 1656–1671 (2016).

81. Lee, J.E. et al. Conversion of Xenopus ectoderm into neurons by NeuroD, a basic helix-loop-helix protein. Science 268, 836–844 (1995).

82. Schwab, M.H. et al. Neuronal basic helix-loop-helix proteins (NEX, neuroD, NDRF): spatiotemporal expression and targeted disruption of the NEX gene in transgenic mice. J Neurosci 18, 1408–1418 (1998).

83. Li, Y., Du, X.F., Liu, C.S., Wen, Z.L. & Du, J.L. Reciprocal regulation between resting microglial dynamics and neuronal activity in vivo. Dev Cell 23, 1189–1202 (2012).

84. Badimon, A. et al. Negative feedback control of neuronal activity by microglia. Nature 586, 417–423 (2020).

85. Gleichmann, M. & Mattson, M.P. Neuronal calcium homeostasis and dysregulation. Antioxid Redox Signal 14, 1261–1273 (2011).

86. Ryczko, D. & Dubuc, R. Dopamine and the Brainstem Locomotor Networks: From Lamprey to Human. Front Neurosci 11, 295 (2017).

87. Yamamoto, K. & Vernier, P. The evolution of dopamine systems in chordates. Front Neuroanat 5, 21 (2011).

88. Vaughan, R.A. & Foster, J.D. Mechanisms of dopamine transporter regulation in normal and disease states. Trends Pharmacol Sci 34, 489–496 (2013).

89. Filippi, A., Mahler, J., Schweitzer, J. & Driever, W. Expression of the paralogous tyrosine hydroxylase encoding genes th1 and th2 reveals the full complement of dopaminergic and noradrenergic neurons in zebrafish larval and juvenile brain. J Comp Neurol 518, 423–438 (2010).

90. Jay, M., De Faveri, F. & McDearmid, J.R. Firing dynamics and modulatory actions of supraspinal dopaminergic neurons during zebrafish locomotor behavior. Curr Biol 25, 435–444 (2015).

91. Yamamoto, K., Ruuskanen, J.O., Wullimann, M.F. & Vernier, P. Two tyrosine hydroxylase genes in vertebrates New dopaminergic territories revealed in the zebrafish brain. Mol Cell Neurosci 43, 394–402 (2010).

92. Filippi, A. et al. Expression and function of nr4a2, lmx1b, and pitx3 in zebrafish dopaminergic and noradrenergic neuronal development. BMC Dev Biol 7, 135 (2007).

93. Wang, Y., Takai, R., Yoshioka, H. & Shirabe, K. Characterization and expression of serotonin transporter genes in zebrafish. Tohoku J Exp Med 208, 267–274 (2006).

94. Guo, S. et al. Development of noradrenergic neurons in the zebrafish hindbrain requires BMP, FGF8, and the homeodomain protein soulless/Phox2a. Neuron 24, 555–566 (1999).

95. Filippi, A., Mueller, T. & Driever, W. vglut2 and gad expression reveal distinct patterns of dual GABAergic versus glutamatergic cotransmitter phenotypes of dopaminergic and noradrenergic neurons in the zebrafish brain. J Comp Neurol 522, 2019–2037 (2014).

96. Manda-Handzlik, A. & Demkow, U. The Brain Entangled: The Contribution of Neutrophil Extracellular Traps to the Diseases of the Central Nervous System. Cells 8 (2019).

97. Tsarouchas, T.M. et al. Dynamic control of proinflammatory cytokines Il-1beta and Tnf-alpha by macrophages in zebrafish spinal cord regeneration. Nat Commun 9, 4670 (2018).

98. Feng, Y., Renshaw, S. & Martin, P. Live imaging of tumor initiation in zebrafish larvae reveals a trophic role for leukocyte-derived PGE(2). Curr Biol 22, 1253–1259 (2012).

99. Tiriac, A. & Feller, M.B. Embryonic neural activity wires the brain. Science 364, 933–934 (2019).

100. Leighton, A.H. & Lohmann, C. The Wiring of Developing Sensory Circuits-From Patterned Spontaneous Activity to Synaptic Plasticity Mechanisms. Front Neural Circuits 10, 71 (2016).

101. Cain, D.W. & Cidlowski, J.A. Immune regulation by glucocorticoids. Nat Rev Immunol 17, 233–247 (2017).

102. Newton, R. Molecular mechanisms of glucocorticoid action: what is important? Thorax 55, 603–613 (2000).

103. Liu, Y.W., Li, S. & Dai, S.S. Neutrophils in traumatic brain injury (TBI): friend or foe? J Neuroinflammation 15, 146 (2018).

104. Hammond, T.R., Robinton, D. & Stevens, B. Microglia and the Brain: Complementary Partners in Development and Disease. Annu Rev Cell Dev Biol 34, 523–544 (2018).

105. Elmore, M.R. et al. Colony-stimulating factor 1 receptor signaling is necessary for microglia viability, unmasking a microglia progenitor cell in the adult brain. Neuron 82, 380–397 (2014).

106. Willis, E.F. et al. Repopulating Microglia Promote Brain Repair in an IL-6-Dependent Manner. Cell 180, 833–846 e816 (2020).

107. Liu, M. et al. Microglia depletion exacerbates acute seizures and hippocampal neuronal degeneration in mouse models of epilepsy. Am J Physiol Cell Physiol 319, C605–C610 (2020).

108. Spangenberg, E.E. & Green, K.N. Inflammation in Alzheimer’s disease: Lessons learned from microglia-depletion models. Brain Behav Immun 61, 1–11 (2017).

109. Dauer, W. & Przedborski, S. Parkinson’s disease: mechanisms and models. Neuron 39, 889–909 (2003).

110. Lam, C.S., Korzh, V. & Strahle, U. Zebrafish embryos are susceptible to the dopaminergic neurotoxin MPTP. Eur J Neurosci 21, 1758–1762 (2005).

111. Madriaga, M.A., McPhee, L.C., Chersa, T., Christie, K.J. & Whelan, P.J. Modulation of locomotor activity by multiple 5-HT and dopaminergic receptor subtypes in the neonatal mouse spinal cord. J Neurophysiol 92, 1566–1576 (2004).

112. Kiehn, O. Locomotor circuits in the mammalian spinal cord. Annu Rev Neurosci 29, 279–306 (2006).

113. Gahtan, E., Tanger, P. & Baier, H. Visual prey capture in larval zebrafish is controlled by identified reticulospinal neurons downstream of the tectum. J Neurosci 25, 9294–9303 (2005).

114. Kimura, Y. et al. Hindbrain V2a neurons in the excitation of spinal locomotor circuits during zebrafish swimming. Curr Biol 23, 843–849 (2013).

115. Reinig, S., Driever, W. & Arrenberg, A.B. The Descending Diencephalic Dopamine System Is Tuned to Sensory Stimuli. Curr Biol 27, 318–333 (2017).

116. Ellett, F., Pase, L., Hayman, J.W., Andrianopoulos, A. & Lieschke, G.J. mpeg1 promoter transgenes direct macrophage-lineage expression in zebrafish. Blood 117, e49–56 (2011).

117. Hall, C., Flores, M.V., Storm, T., Crosier, K. & Crosier, P. The zebrafish lysozyme C promoter drives myeloid-specific expression in transgenic fish. BMC Dev Biol 7, 42 (2007).

118. Peri, F. & Nusslein-Volhard, C. Live imaging of neuronal degradation by microglia reveals a role for v0-ATPase a1 in phagosomal fusion in vivo. Cell 133, 916–927 (2008).

119. Kimmel, C.B., Ballard, W.W., Kimmel, S.R., Ullmann, B. & Schilling, T.F. Stages of embryonic development of the zebrafish. Dev Dyn 203, 253–310 (1995).

120. Earley, A.M., Dixon, C.T. & Shiau, C.E. Genetic analysis of zebrafish homologs of human FOXQ1, foxq1a and foxq1b, in innate immune cell development and bacterial host response. PLoS One 13, e0194207 (2018).

121. Contreras-Lopez, O., Moyano, T.C., Soto, D.C. & Gutierrez, R.A. Step-by-Step Construction of Gene Co-expression Networks from High-Throughput Arabidopsis RNA Sequencing Data. Methods Mol Biol 1761, 275–301 (2018).

